# Microbial ecology and site characteristics underlie differences in salinity-methane relationships in coastal wetlands

**DOI:** 10.1101/2024.04.02.587477

**Authors:** Clifton P. Bueno de Mesquita, Wyatt H. Hartman, Marcelo Ardón, Emily S. Bernhardt, Scott C. Neubauer, Nathaniel B. Weston, Susannah G. Tringe

## Abstract

Methane (CH_4_) is a potent greenhouse gas emitted by archaea in anaerobic environments such as wetland soils. Tidal freshwater wetlands are predicted to become increasingly saline as sea levels rise due to climate change. Previous work has shown that increases in salinity generally decrease CH_4_ emissions, but with considerable variation, including instances where salinization increased CH_4_ flux. We measured microbial community composition, biogeochemistry, and CH_4_ flux from field samples and lab experiments from four different sites across a wide geographic range. We sought to assess how site differences and microbial ecology affect how CH_4_ emissions are influenced by salinization. CH_4_ flux was generally, but not always, positively correlated with CO_2_ flux, soil carbon, ammonium, phosphate, and pH. Methanogen guilds were positively correlated with CH_4_ flux across all sites, while methanotroph guilds were both positively and negatively correlated with CH_4_ depending on site. There was mixed support for negative relationships between CH_4_ fluxes and concentrations of alternative electron acceptors and abundances of taxa that reduce them. CH_4_/salinity relationships ranged from negative, to neutral, to positive and appeared to be influenced by site characteristics such as pH and plant composition, which also likely contributed to site differences in microbial communities. The activity of site-specific microbes that may respond differently to low-level salinity increases is likely an important driver of CH_4_/salinity relationships. Our results suggest several factors that make it difficult to generalize CH_4_/salinity relationships and highlight the need for paired microbial and flux measurements across a broader range of sites.

## Introduction

The most recent report by the Intergovernmental Panel on Climate Change stated that increases in well mixed greenhouse gases such as carbon dioxide (CO_2_), methane (CH_4_), and nitrous oxide (N_2_O) have contributed 1.0 - 2.0°C of warming to Earth’s climate in the past century (1). After CO_2_, CH_4_ is the largest contributor, with a global warming potential 84-86 times that of CO_2_ over a 20-year timeframe. CH_4_ concentrations have increased from 675 ppb in the 1700s to 1866 ppb in 2019, after being stable for most of the previous 800 years (1, 2).

Wetlands are the largest natural source of CH_4_; the methanogenic activity of anaerobic archaea in wetland soils contributes approximately 30% of all methane emissions globally (3). It is thus important to understand the drivers of wetland CH_4_ emissions, especially given that there has been much recent discussion of the potential for coastal and estuarine wetlands to sequester vast quantities of carbon in what has been referred to as “blue carbon” (4–7). However, any increases in carbon (C) storage could potentially be offset by greenhouse gas emissions (8–10), which warrants further study and quantification.

Another topic in need of further research is how minor increases in salinity will affect greenhouse gas emissions, especially given that two principal factors driving coastal and estuarine wetland salinization - drought and sea level rise - are predicted to increase in the future with ongoing climate change (11). Such salinization causes former tidal freshwater wetlands to become oligohaline (> 0.5 to 5 ppt). Globally, sea levels have risen 20 cm since 1901 and are currently (2006–2018) rising at a rate of 3.7 mm per year, as a result of both melting ice and thermal expansion (1). Droughts have increased in frequency and severity in many parts of the world, which decreases freshwater inputs into estuarine and coastal ecosystems, leading to low-level increases in salinity (1, 12). Several other anthropogenic factors such as water management can also contribute to low-level salinization, which, in turn, precipitates ecosystem changes including loss of biodiversity and ecosystem services (13).

Salinity is a key environmental variable that affects plant, bacterial, archaeal, fungal, and zooplankton community composition and productivity, as well as biogeochemical processes (14–18). Only certain organisms in the tree of life have evolved to function at elevated salt concentrations, either by using compatible solutes to keep salt out or by having acidic proteomes and special ion pumps to function with high intracellular salt concentrations (19, 20). Effects of salinity on plant communities have been well documented, with drastic observed compositional changes along salinity gradients, including low-level, or oligohaline, salinities (21). High salinities are toxic to plants adapted to freshwater, while plants adapted to saltwater are outcompeted by freshwater-adapted plants in freshwater environments, and are therefore only found in saltwater environments (14). Salinity has been found to be a principal variable structuring microbial communities at global, regional, and landscape scales (16, 22–24). At the chemical level, salinity also affects the solubility of dissolved organic carbon and other compounds, as well as the sorption of inorganic ions (25–27). While oligohaline salinities are high enough to elicit responses in microbial communities (28, 29), work done across broader salinity gradients including mesohaline (5-18 ppt) and polyhaline (18-30 ppt) areas suggests that the effects of salinization will likely be dependent on the magnitude of salinization (23, 30, 31). This may in turn affect the relationship between salinity and methane emissions, which has been shown to be non-linear (23).

Theory predicts that CH_4_ emissions will decrease with increased seawater influence as the concurrent increase in sulfate from seawater will promote sulfate-reducing organisms that can outcompete methanogens for shared resources, such as acetate and hydrogen (32–36). Such a decrease, in fact a log-linear decrease in CH_4_ emissions with salinity (including oligohaline salinities up to seawater salinities), is indeed what has been found in several studies and meta-analyses (37–40). However, a summary of laboratory microcosm experiments that tested salinity-methane relationships reported 8 negative relationships, 2 *positive* relationships, and 1 neutral relationship (41). Similar variation has been observed for CO_2_ emissions (40). Such variation in relationships has been attributed in part to hydrological setting (41, 42) and soil characteristics (43), but the roles of site context/history, including legacy effects of agriculture and fertilizer use (44), as well as microbial ecology – relations of microbes to one another and to their physical surroundings - are two other main factors that could contribute to such discrepancies and are in need of further research. Wetlands with a history of agriculture use, and consequently fertilizer application, are characterized by what are known as biogeochemical legacies, including altered carbon and nutrient pools, such as legacy nitrogen and phosphorus that can remain in soils for decades after the cessation of inputs (44). Other important site legacies include the history of inundation and prior exposure to salinity (45).

CH_4_ cycling is driven by the activity of, and interactions among, multiple distinct functional groups of microorganisms, including complex carbon degraders, fermenters, sulfate reducers, iron reducers, ammonia oxidizers, denitrifiers, methanogens, and methanotrophs (46–48). These groups encompass functionally and taxonomically diverse organisms. Complex carbon degraders and fermenters are crucial players in the decomposition process, breaking down larger, more recalcitrant carbon compounds such as lignin and cellulose into fatty acids and eventually acetate, H_2_ and CO_2_ that fuel methanogenesis (47). While sulfate reducers can compete with methanogens, they can also be syntrophic with methanogens, as certain taxa produce acetate and hydrogen which would then fuel methanogenesis from those substrates.

Methanogens can perform one of four different methanogenesis pathways, including acetoclastic (CH_4_ produced from acetate), hydrogenotrophic (CH_4_ produced from hydrogen and CO_2_ or other electron acceptors such as carbon monoxide or alcohols), methyl-dismutation (i.e., methylotrophic, CH_4_ produced from methylated compounds), and methyl-reduction (CH_4_ produced from methylated compounds and hydrogen) (49, 50). Most methyl-dismutation pathways are likely not affected by competition with sulfate reducers (51–54). Methanotrophs include anaerobic archaea and both anaerobic and aerobic bacteria (55, 56), the latter of which have been divided into separate classes (I, II, IIa) (57). Furthermore, prior work has suggested key interactions between the methane and nitrogen cycles, and among the taxa involved, particularly methanotrophs, ammonia oxidizers (AO) and nitrite oxidizing bacteria (NOB) (48). Nitrogen can stimulate CH_4_ oxidation in N limited conditions, while excess ammonia and nitrite can inhibit CH_4_ oxidation (58–60). Such diversity of microorganisms may lead to contrasting responses or a lack of response to low salinity, depending on the direct effects of salinity on these taxa, the indirect effects of other environmental variables on these taxa, the effects of interactions among these taxa, and the overall functional redundancy of the microbial community (57, 61, 62).

In this study we combine microbial and biogeochemical data from field and laboratory experiments in four different sites to assess how salinization affects microbial communities and biogeochemistry, including greenhouse gas fluxes. These data encompass tidal freshwater marshes, oligohaline wetlands, a freshwater forested wetland, control microcosms and microcosms amended with artificial seawater (ASW), field plots amended with artificial seawater, and tidal freshwater soils transplanted to an oligohaline wetland. With respect to microbial ecology, we hypothesized that (1) low salinity would reduce alpha-diversity within each site due to the direct effects of NaCl on freshwater-adapted taxa, and (2) broad site geographic, hydrologic, plant, and biogeochemical differences would lead to significant differences in microbial beta-diversity across the four geographic locations due to the effects of those variables on microorganisms. We also predicted that some of the same taxa that are particularly sensitive to or adapted to salt would respond consistently to natural and experimental salinity gradients. With respect to relationships with CH_4_ flux, we hypothesized that (1) a greater relative abundance of methanogens would fuel more methanogenesis and a lower relative abundance of methanotrophs results in more CH_4_ emitted to the atmosphere, (2) increased availability of alternative electron acceptors would suppress methanogenesis due to competition, and (3) greater overall decomposition rates would fuel more methanogenesis by providing methanogenic substrates. More specifically, we predicted that (1) all methanogen guilds and the methanogen: methanotroph ratio would be positively correlated with CH_4_ flux, while all methanotroph guilds would be negatively correlated with CH_4_ flux, (2) CH_4_ flux would be negatively correlated with salinity, concentrations of alternative electron acceptors (nitrate, sulfate, iron, manganese), and abundances of taxa that reduce them (iron reducing bacteria and sulfate reducing bacteria); and (3) CH_4_ flux would be positively correlated with correlates of decomposition, including CO_2_ flux, organic carbon, ammonium, phosphate, pH, and abundance of hypothesized complex C degraders (Actinobacteriota, Firmicutes). While we did not directly measure decomposition rates, the literature shows that decomposition rates are positively correlated with carbon availability (63), as well as nutrient availability, including nitrogen and phosphorus (64, 65). Research has also shown that decomposition rates are higher at more neutral pH compared to acidic pH (66). Firmicutes and Actinobacteriota were shown to harbor complex carbon degradation genes in wetlands (23), so we also include them in this prediction.

## Methods

We synthesized data from five research projects from four separate sites, chosen because they included measurements of methane fluxes in freshwater and oligohaline conditions, either due to natural variation or to field or lab manipulation, and had retained soil samples adequate for DNA extraction and sequencing. The projects include field sampling across a natural salinity gradient, field manipulations of salinity with either seawater addition or transplanting, and laboratory microcosm incubation experiments in which soils were brought back to the laboratory to receive artificial seawater (ASW) additions for at least 12 weeks. Broadly, the four sites are the San Francisco Bay Estuary in California, USA (“SF”) (23), the Delaware River Estuary in New Jersey, USA (“DE”) (30), the Alligator River Estuary in North Carolina, USA (“Alli”) (26, 67), and the Waccamaw River Estuary in South Carolina, USA (“Wacc”) (68) (Table 1, Figure S1). These estuaries contain vast expanses of wetlands; for example, there are ∼59,000 ha of wetlands in the San Francisco Estuary and ∼445,000 ha and of wetlands in the Delaware Estuary. From all of the potential data available, we selected only samples with freshwater (< 1 ppt) or oligohaline (1 - 5 ppt) salinity classes, as this is the most important salinity class to understand regarding seawater intrusion into freshwater wetlands and tidal marshes (Table 1).

All soil samples were covered with water and come from similar depth classes within the 0-5 cm and 5-15 cm range. All soil samples were exposed to increased salinity for at least 12 weeks, but the length of exposure varied among studies, including decades (San Francisco), 3 years (Waccamaw), 1 year (Delaware field), and 12 weeks (Delaware lab, Alligator lab). CH_4_ flux was measured at all sites; however, the surface area, volume, and depth of the sample upon which measurements were taken were different at each site, and some sites used a gas chromatograph while a portable analyzer was used at San Francisco (Table 1). Thus, methane flux data are comparable among treatments in the same site but are not directly comparable among the sites (Table 1). Additionally, CO_2_ flux was measured at San Francisco, Waccamaw, and Alligator, and N_2_O flux was also measured at Delaware and Alligator. Suites of other soil and porewater variables were measured, and this varied among the sites and experiments as described below. Variables measured at only one site were included in the analysis if similar variables were measured at other sites (e.g., soil organic matter, dissolved organic carbon).

### Sampling and Measurements

In San Francisco, soils were collected from 2 freshwater and 2 oligohaline wetland complexes in the delta formed by the Sacramento and San Joaquin rivers as they empty into the San Francisco Bay. Each salinity class included one unaltered reference wetland and one wetland restored from agricultural use as either cropland or pasture. At each wetland, three cores were taken with a split core auger with an airtight plastic sleeve with a 5 cm diameter and 15 cm depth. Each core in the plastic sleeve was immediately capped on both ends to maintain an anaerobic environment. The core was then placed into a 2 L Mason jar and the top cap was removed to quantify CH_4_ and CO_2_ flux over the course of 5 minutes on an ultraportable greenhouse gas analyzer (Los Gatos Research, San Jose, CA, USA). An additional core adjacent to each initial core was taken, split into the 0-5 cm depth segment and 5-15 cm depth segment, and aliquoted for DNA sequencing (stored at −20°C) and biogeochemical analyses of soil and porewater at the UC Davis Analytical Laboratory. Measured variables included: porewater dissolved organic carbon (DOC) and pH, soil C, nitrogen (N), C:N, ammonium (NH_4_^+^), and Olsen phosphate (PO_4_^3-^), and both porewater and soil nitrate (NO_4_^-^), sulfate (SO_4_^2-^), iron (Fe), and manganese (Mn). Samples from San Francisco are all unmanipulated field samples (n = 72).

At Waccamaw, a field experiment was established in the 0.9 ha Brookgreen Gardens tidal freshwater marsh (33°31.50′ N, 79°5.51′ W), adjacent to Springfield Creek, a tidal tributary of the Waccamaw River (68). The site is flooded 10-30 cm during most high tides. Five replicate plots, at least 3 m apart from each other, received 40 L of either freshwater or seawater additions every 3-4 days over the course of three years. Seawater additions were from salt marsh tidal creek water from the flow-through seawater system at the Baruch Marine Field Laboratory (BMFL) that was diluted with freshwater to a salinity of 10.2 before being added to the marsh.

Freshwater additions were from a 180-m-deep groundwater well at the BMFL (salinity = 0.5 ± 0.04). Plot-scale exchanges of CO_2_ and CH_4_ were measured with large, transparent, temperature-controlled chambers 0.37 m^2^ ×1.22 m height. Air within each chamber was stirred with four fans while pumps circulated air between the chambers and a LI-COR LI7000 CO_2_/H2O analyzer (LI-COR Biosciences, Lincoln, NE, USA). To determine CH_4_ fluxes, air samples were collected from each chamber roughly every 5–7 min, stored in gas tight Hungate tubes, and analyzed for CH_4_ concentration in the laboratory using a flame ionization detector on a Shimadzu GC14A gas chromatograph (Shimadzu Scientific Instruments, Columbia, MD, USA). Soils were collected at the end of the experiment by collecting one core of 6.4 cm diameter and 10 cm depth from each plot. Other measured variables included soil organic matter (SOM), C, N, and C:N, and soil oxygen demand (SOD). Samples from Waccamaw are all manipulated field samples (n = 30).

At Delaware, data come from two separate projects - a field transplant experiment and a laboratory incubation experiment. For the Delaware transplant experiment, large (29 cm diameter and approximately 30 cm depth) intact cores of tidal freshwater marsh (Rancocas Creek) soil and vegetation were transplanted into an oligohaline wetland (Salem) and a separate tidal freshwater marsh (Raccoon Creek) to serve as a control. Tidal freshwater marsh plant communities were dominated by *Peltandra virginica* and species of *Bidens*, *Amaranthus*, and *Polygonum*, while the oligohaline site was dominated by *Zizania aquatica* (wild rice).

Transplants were done at water level as well as 40 cm below water level, but only the 40 cm flooded samples are included here, to more closely match the Delaware laboratory data (see below) and the Waccamaw field experiment (see above). Soil for DNA sequencing was taken from 0-4 cm and 10-14 cm depth. Greenhouse gas measurements were taken by placing covered (dark) acrylic chambers of 43 L volume over the soil cores. Carbon dioxide production inside the chamber was measured with a PPSystems infrared gas analyzer for several minutes. Methane exchange was measured by taking gas samples at 0, 10, and 20 minutes with a 60 mL syringe, injecting into 20 mL evacuated vials, and measuring CH_4_ on an Agilent 6890N gas chromatograph with a flame ionization detector. Other measured variables here included porewater NH_4_^+^, PO_4_^3-^, SO_4_^2-^, and Fe^2+^.

For the Delaware laboratory microcosm incubation experiment, soils from the Woodbury Creek tidal freshwater marsh were collected to a depth of 25 cm with 10 cm diameter polyvinyl chloride tubes and sealed with gas- and water-tight caps (69). Holes were drilled in the core barrel just above the soil surface, and then cores were placed into separate 100 L tidal tanks in the dark at 20°C. The tidal tanks simulate the tidal cycle with 6 h of air exposure followed by 6 h of inundation. Tanks were filled with artificial freshwater that mimicked the freshwater chemistry of the Delaware River. After a 2-week pre-incubation period, water in one tank was replaced with artificial seawater with a salinity of 4.95 ppt. Both tanks were changed at least once weekly after the initial 2-week period. Cores were incubated for 12 weeks after which the 0-4 cm and 10-14 cm depth portions were sampled and stored at −20°C for DNA sequencing, with an aliquot sent for porewater and soil biogeochemical measurements which included Cl^-^, SO_4_^2-^, PO_4_^3-^, DOC, NO_x_, and acetate. Gas flux was measured by fitting a 1.2 L gas-tight cap onto the core. An initial headspace sample (3 mL) for CH_4_ analysis was obtained with a gas-tight syringe; then final CH_4_ samples were obtained after approximately 1 h. CH_4_ samples were analyzed immediately by flame ionization detection gas chromatography (Agilent 6890 N with Porapak Q column). The wetlands that were sampled in both experiments have no known prior history of agricultural use but are affected by runoff from surrounding agricultural and urban areas. Sample sizes from Delaware are n = 8 for the transplant experiment and n = 8 for the laboratory experiment (Table 1).

At Alligator, soils were collected from the Timberlake Observatory for Wetland Restoration, a forested freshwater wetland restored in 2004 from prior use as a corn field that is now used as a research site. The site had not experienced saltwater incursion for at least 20 years prior to sampling. The site where the soils were collected is characterized by Eutric Histosol soils and Atlantic white cedar vegetation (26). A laboratory incubation was started with intact soil cores 2.5 cm in diameter and 30 cm deep. Alligator microcosms received either deionized freshwater (control), artificial seawater, artificial seawater without sulfate, or sulfate. In this analysis we only included the controls and artificial seawater additions, as in the Delaware experiment. To measure CO_2_, CH_4_, and N_2_O, cores were fitted with a gas tight lid with a Swagelok brass sampling port with a rubber septum (0.6 cm). Headspace gas samples were collected immediately and after 1 h into evacuated 8 ml gas vials. Gases were quantified on a Shimadzu 17A gas chromatograph with electron capture detector (ECD), flame ionization detector (FID), and methanizer (Shimadzu Scientific Instruments, Columbia, MD, USA). As in Delaware, the experiment proceeded for three months, after which soils from 0-5 cm and 10-15 cm depths were collected, a portion stored at −20°C for DNA sequencing, and another aliquot analyzed for porewater biogeochemistry. Measured variables at Alligator were porewater NO_3_^-^, SO_4_^2-^, total organic carbon (TOC), dissolved organic nitrogen (DON), dissolved inorganic nitrogen (DIN), total nitrogen (TN), NH_4_^+^, PO_4_^3-^, and soil pH, %C, %N and C:N.

### Microbial sequencing and analysis

Soils were frozen at −20°C until DNA extraction. DNA was extracted from 0.3 g of soil with a Qiagen DNeasy PowerSoil kit following the manufacturer’s instructions. PCR was then used to amplify the V4 region of the 16S rRNA gene, following the standard methods of the U.S. Department of Energy Joint Genome Institute (70). DNA was sequenced on a MiSeq 2000 (Illumina Inc., CA, USA) with paired-end 150 base pair chemistry. Raw data were processed with the iTagger pipeline to quality-filter reads, dereplicate sequences, and cluster sequences into operational taxonomic units (OTUs) at 97% sequence similarity (70). Taxonomy was assigned using the ‘assignTaxonomy’ function in the *dada2* R package (71), with the SILVA 138.1 taxonomic database (72). The *mctoolsr* R package (73) was used to remove chloroplast and mitochondrial DNA and any taxa that were not assigned at least to Bacteria or Archaea at the domain level. We used known taxonomy-function relationships to assign functional guilds of interest for anaerobic biogeochemistry (23), using the “Get_16S_guilds_alt” function in a publicly available R script (https://github.com/cliffbueno/SF_microbe_methane/blob/main/modules/AssignGuilds.R).

Sequencing depth was 131845 ± 3373 SE sequences per sample, and data were rarefied to 31,264 sequences per sample. Only one sample from San Francisco with very few reads (1296) was dropped. The final sample size analyzed was 133 (Table 1). Sequences were deposited to NCBI GenBank with BioProject ID PRJNA1004999.

### Statistical analysis

The number of OTUs observed per sample and Shannon diversity were used as microbial alpha-diversity metrics. The effects of site, salinity class (freshwater or oligohaline), and depth (surface 0-5 cm range versus deeper 5-15 cm range) on alpha-diversity were tested with ANOVA followed by Tukey’s post hoc. Microbial community composition was assessed by calculating a Bray-Curtis dissimilarity matrix on square-root transformed abundances, and performing a PERMANOVA test implemented with the ‘adonis2’ function in the *vegan* R package (74). Within-group multivariate homogeneity of dispersion was tested with PERMDISP implemented with the ‘betadisper’ function in *vegan*. To compare a presence/absence-based metric with the Bray-Curtis metric, we also calculated the Jaccard dissimilarity metric with *vegan*. Unique and overlapping taxa in freshwater and oligohaline salinity classes were calculated with *mctoolsr*. Differences in relative abundances of taxa among the salinity classes were tested with Wilcoxon tests. Indicator species analysis to identify taxa associated with freshwater or oligohaline salinities was performed with the ‘multipatt’ function in the *indicspecies* R package (75), with the ‘r.g’ species-site group association function. A phylogenetic tree was built by aligning sequences with MUSCLE (76) and then building a tree with fasttree (77), both of which were implemented in QIIME (78). Nearest taxonomic index (NTI) was calculated for each sample using the ‘NTI.p’ function in the *iCAMP* R package (79). NTI is a standardized measure of the phylogenetic distance to the nearest taxon for each taxon in a sample. NTI values > 2 or < −2 suggest deterministic processes govern community assembly while NTI values between −2 and 2 suggest dominance of stochastic processes (80). Models of environmental predictors of community composition were tested with distance-based redundancy analysis (dbRDA), implemented with the ‘capscale’ function in *vegan*; the best combination of predictors was selected using backward model selection with the ‘ordistep’ function in *vegan*. Communities were visualized with principal coordinates analysis (PCoA), with environmental vectors fit with the ‘envfit’ function in *vegan*. Spearman correlations between CH_4_ flux and chemical variables and certain microbial guild or taxa abundances were calculated and p-values corrected with false discovery rate (FDR). All figures were made with either the *ggplot2* (81) or *pheatmap* (82) R packages. All analyses were performed with R version 4.0.2 (83).

## Results

All wetland soils measured in this study emitted a net flux of CH_4_ to the atmosphere, the amount of which varied by at least three orders of magnitude (Figure 1). Among the five datasets, there were discrepancies in the relationship between salinity and CH_4_ fluxes, with 2 positive relationships, 1 negative relationship, and 2 neutral relationships (Figure 1).

**Figure 1.**
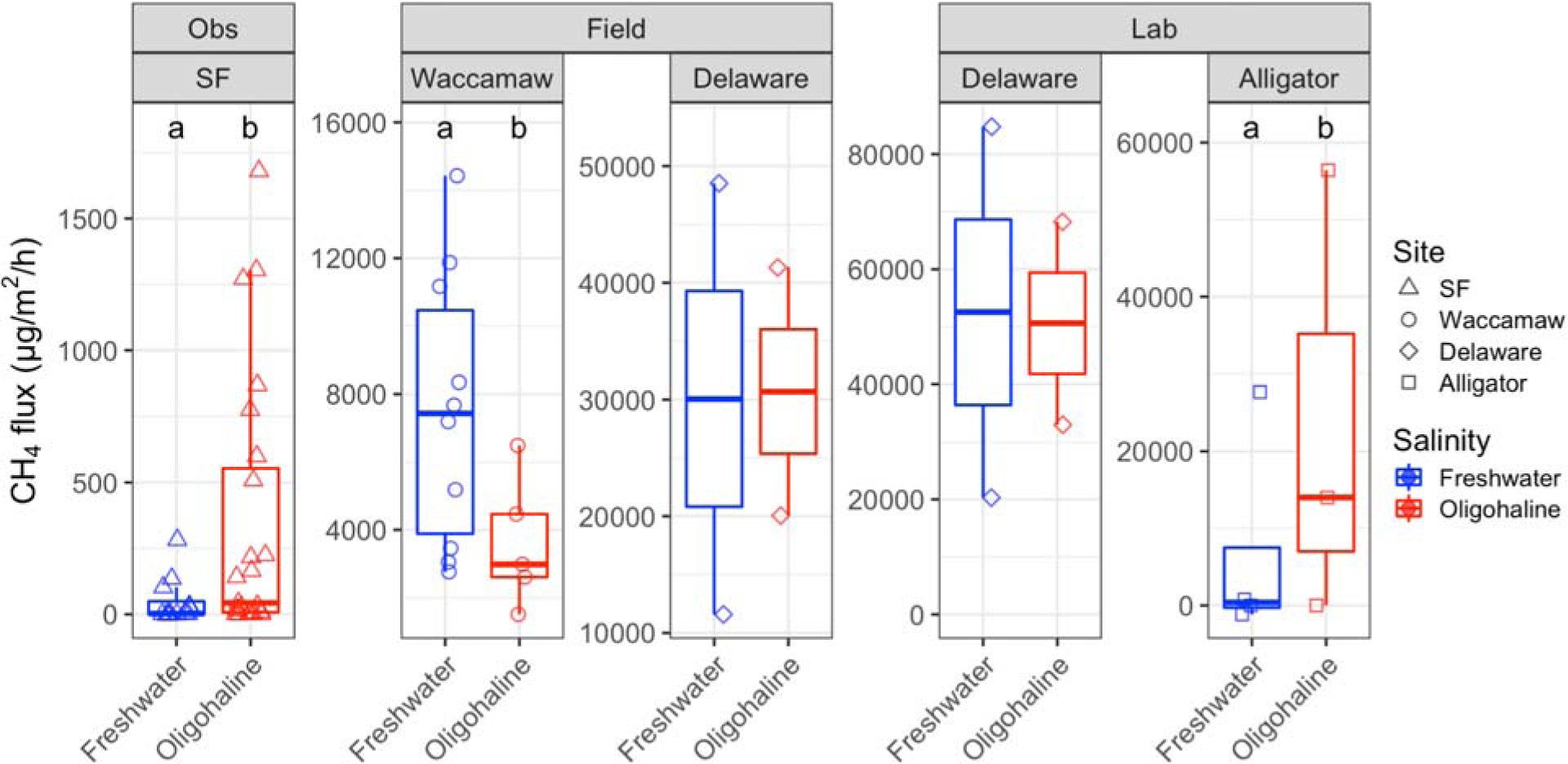
Methane (CH_4_) flux from the observational study (Obs), both field experiments, and both lab experiments. Note the difference in y-axis scales among panels and that each study used a different method for measuring CH_4_ flux, so absolute values of fluxes are not comparable across panels, only within panels. Sample sizes per study are 36 for San Francisco, 15 for Waccamaw, 4 for Delaware field, 4 for Delaware lab, and 15 for Alligator (Table 1). Different letters represent significant differences (t-test, p < 0.05). Data are only shown for cores with microbial data. Data from the full experiments at Waccamaw and Alligator are consistent with results shown here. Data from the full Delaware lab experiment show increased CH_4_ flux in the oligohaline treatment.

Microbial alpha diversity metrics, including OTU richness and Shannon diversity, differed significantly among sites, salinity classes, and depths (Figure 2, Table 2). In two sites, San Francisco and Waccamaw, richness in oligohaline samples was significantly lower than richness in freshwater samples, as was Shannon diversity in San Francisco and Alligator (Tukey HSD p < 0.05, Figure 2). Alpha diversity was not significantly affected by salinity class in the Delaware field or lab experiments.

**Figure 2.**
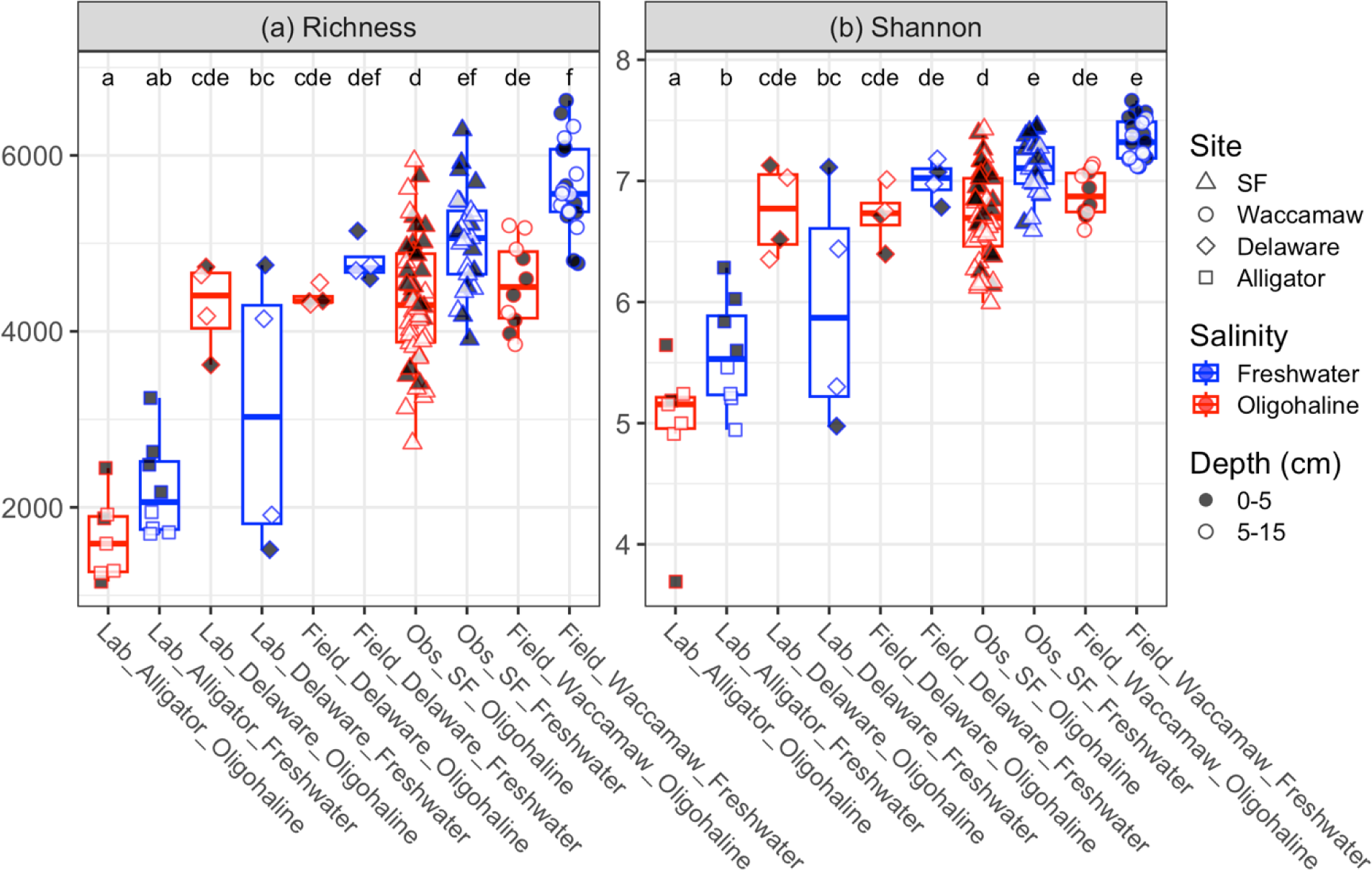
OTU richness (a) and Shannon diversity (b) of bacterial and archaeal soil communities across four different sites, two salinity classes, and two depth ranges. DE = Delaware, SF = San Francisco. Sample sizes per study are 36 for SF, 15 for Waccamaw, 4 for Delaware field, 4 for Delaware lab, and 15 for Alligator (Table 1). Different letters represent significant pairwise comparisons within each panel (Tukey posthoc, p < 0.05). The x-axis is sorted by increasing ASV richness by study (combined freshwater and oligohaline data for each study), with freshwater on the left and oligohaline on the right, for each study. Study type (lab, field, or observational (Obs)) is stated in the x-axis labels.

Microbial community composition was primarily driven by site and secondarily by salinity, pH, depth, and suites of other environmental factors that were unique to each site (Figure 3, Table 2). When analyzing all of the data together, samples from each site, including all depths, salinities, and environmental gradients within each site, clustered together, demonstrating a primary influence of site. Communities associated with higher salinities were also associated with lower CH_4_ emissions (Figure 3a, ‘envfit’ p < 0.05).

**Figure 3.**
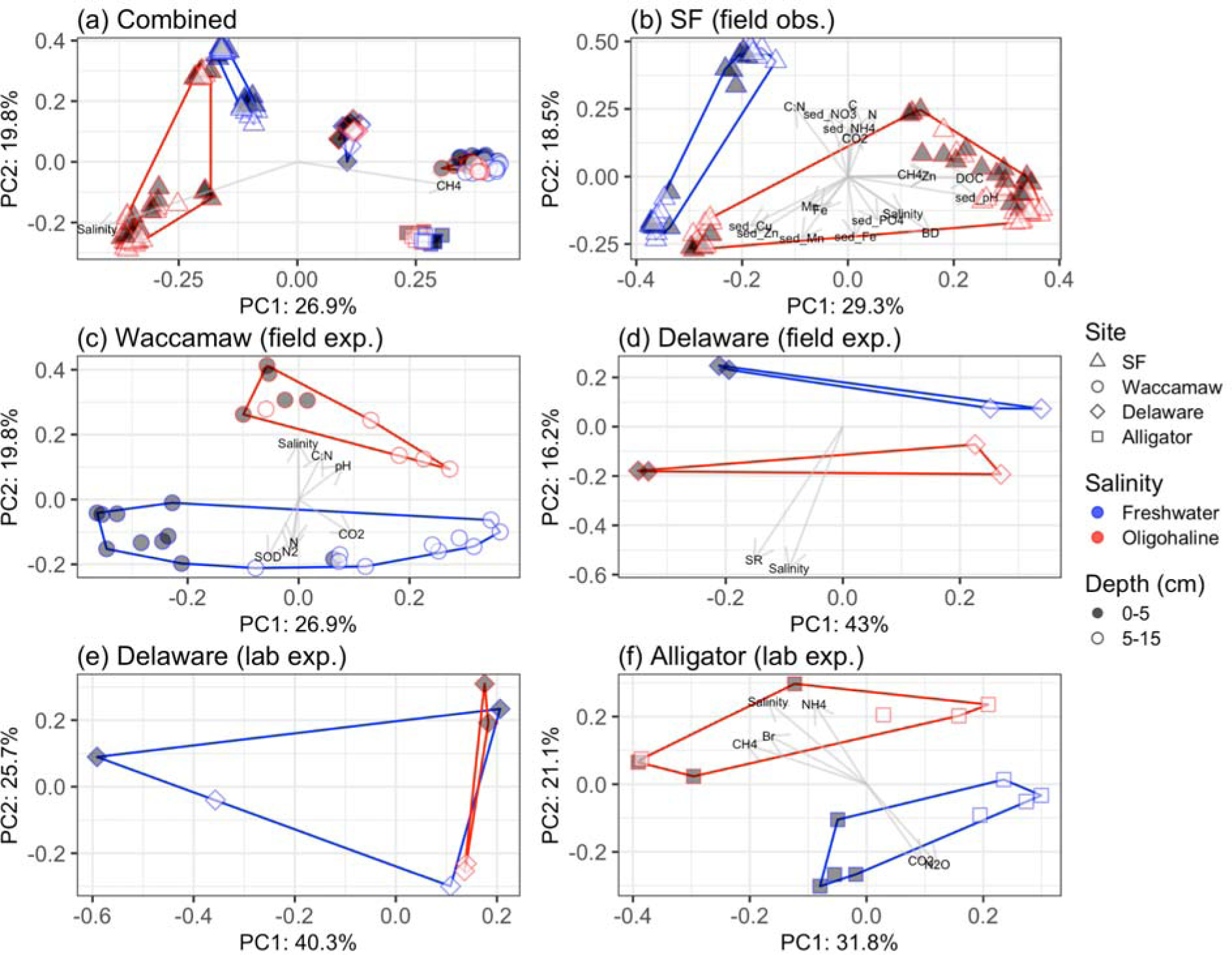
Principal coordinates analysis of Bray-Curtis dissimilarities at the OTU level for bacterial and archaeal soil communities for all data from all four estuaries (a), San Francisco Bay field salinity gradient (b), Delaware River estuary field experiment (c), Waccamaw River estuary field experiment (d), Delaware River laboratory incubation experiment (e), and Alligator River estuary laboratory incubation experiment (f). Sample sizes per study 36 for SF, 15 for Waccamaw, 4 for DE field, 4 for DE lab, and 15 for Alligator (Table 1). Note that salinity and CH_4_ were the only two continuous variables measured in all five studies (shown as vectors in panel a).

To further assess the role of environmental factors within sites, tests were also performed for each site/experiment separately. In San Francisco, a field study with the highest sample size, many environmental variables, including CO_2_ flux, CH_4_ flux, salinity, bulk density, soil C, N, C:N, NO_3_^-^, NH_4_^+^, pH, Cu, Zn, Mn, Fe, PO_4_^3-^, and porewater DOC, and Zn, were correlated with community composition (Figure 3b). Stepwise redundancy analysis model selection suggested that bulk density, soil Zn, pH, Cl^-^, C:N, and Mn, and porewater sulfate were the most important variables driving microbial community composition. In the Waccamaw field experiment, results were similar; samples in plots that received seawater additions and thus became oligohaline, and with higher C:N ratios and pH, were significantly different from control freshwater samples, which had higher CO_2_ flux, soil N concentrations, net N_2_ emissions, and SOD. There was also a significant effect of depth (Figure 3c). The best model predicting community composition contained SOD and CO_2_ flux as variables. In the Delaware field transplant experiment, oligohaline samples, associated with increased salinity and sulfate reduction, were significantly different from freshwater samples. The best model predicting community composition only contained salinity as a predictor variable. Depth was also a particularly important factor at this site; samples from the same depth but different salinities were more similar than vice versa (Figure 3d). Laboratory incubations with Delaware wetland soils did not cleanly cluster by depth and salinity class as the other experiments did, due to a high degree of variability in the control (freshwater salinity) samples and a lower sample size (Figure 3e). Community dispersion was not homogenous among the two salinity classes (PERMDISP, p < 0.05), indicating this higher degree of variability within the control group. On the other hand, the laboratory incubations with Alligator soils did cluster by salinity class and depth. Samples that received ASW addition making them oligohaline were associated with higher NH_4_^+^, Br^-^, and CH_4_ while controls were associated with higher CO_2_ and N_2_O fluxes (Figure 3f). SO_4_^2-^, PO_4_^3-^, and CH_4_ flux were the three variables selected to best predict microbial community composition in stepwise redundancy analysis.

Across the whole dataset, the most abundant bacterial phyla were Proteobacteria (mean = 21%), Acidobacteriota (10%), Chloroflexi (8%), Bacteroidota (8%), Desulfobacterota (7%), Firmicutes (7%), Actinobacteriota (6%), Verrucomicrobiota (5%), Planctomycetota (5%), Nitrospirota (3%), Myxococcota (3%), and Crenarchaeota (2%) (Figure 4). Within sites, many of these phyla differed significantly between freshwater and oligohaline samples (Figure 4), with some consistent responses detected in Verrucomicrobiota (negative response in 3/4 sites), Proteobacteria and Myxococcota (negative response in 2/4 sites), and Firmicutes and Chloroflexi (positive response in 2/4 sites) (Figure 4). There were also some key differences in dominant phyla among the sites, with Alligator characterized by a much higher percentage of Acidobacteriota and Firmicutes than the other sites.

**Figure 4.**
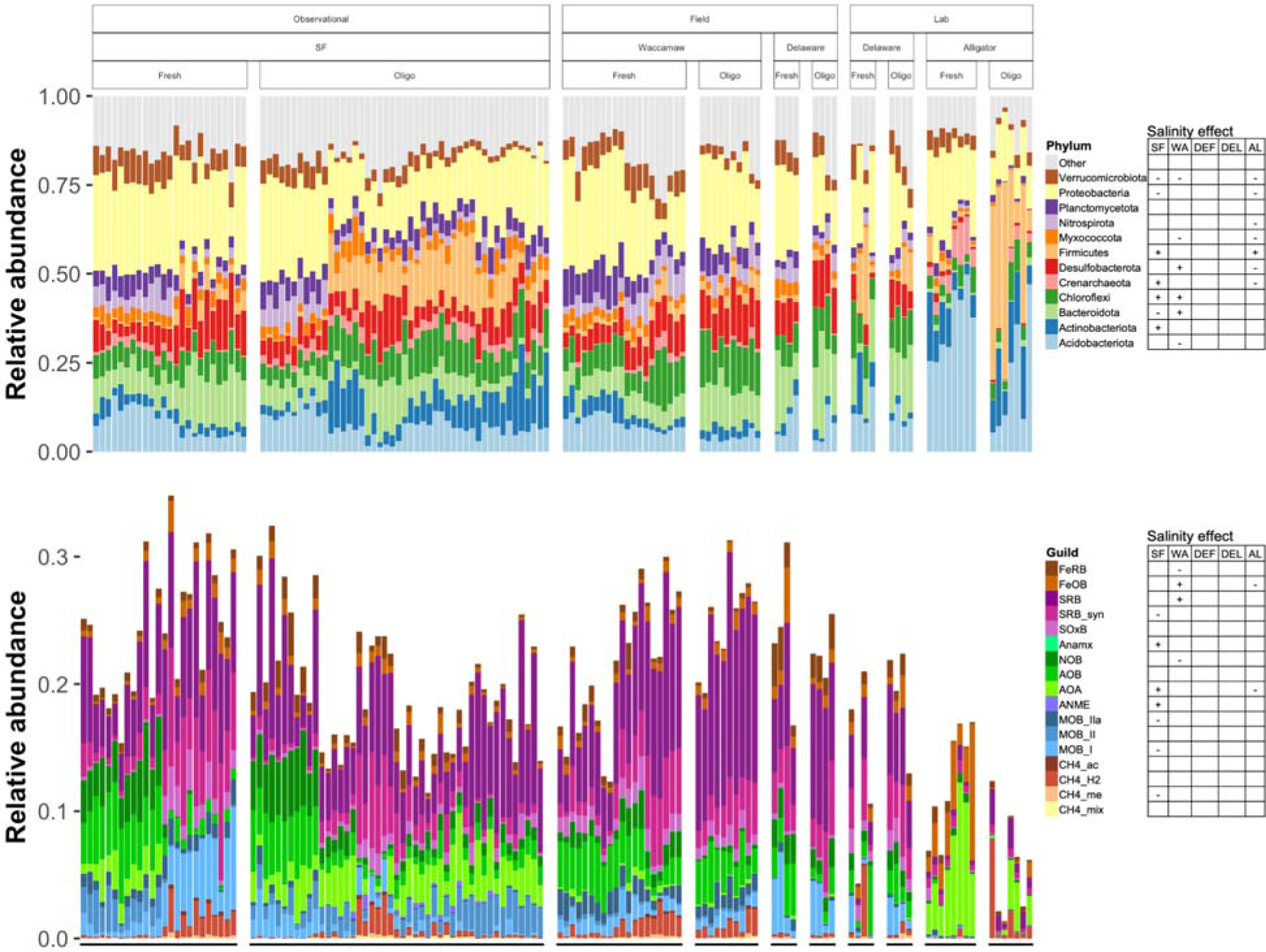
Relative abundance of the top 12 phyla (top panel) and functional guilds (bottom panel), and whether there is a significant positive (+) or negative (-) effect of salinity on the phylum or guild. Relative abundances in the top panel sum to 1, as all other taxa are included in the “Other” category. Relative abundances in the bottom panel are aggregated counts of OTUs that are assigned to those guilds. DEF = Delaware field experiment; DEL = Delaware lab experiment. Each column represents an individual sample, but sample IDs are omitted from the x-axis label for clarity.

The most abundant microbial functional guilds across all samples were sulfate reducing bacteria (mean = 6%), syntrophic sulfate reducing bacteria (2%), ammonia oxidizing bacteria (2%), ammonia oxidizing archaea (2%), nitrite oxidizing bacteria (1%), and class I methane oxidizing bacteria (1%) (Figure 4). All other guilds made up less than 1% of the community on average across all sites, although in some individual samples they had greater relative abundances. Across all samples, methanogens averaged 0.8% relative abundance and methanotrophs averaged 2.7% relative abundance. Most guilds varied significantly among sites and within sites some were differentially abundant between freshwater and oligohaline samples. However, unlike some of the dominant phyla, there were no consistent responses to salinity among guilds across multiple sites (Figure 4).

There were 220 methanogen OTUs: 146 hydrogenotrophic OTUs, 29 acetoclastic OTUs, 35 methyl-reducing OTUs, and 10 mixotrophic OTUs. 44 methanogen OTUs were found in at least two sites. Methanogen community composition varied among the 4 sites (Figure S3). Hydrogenotrophs were the most abundant methanogen guild across the whole dataset, followed by methyl reducers, acetoclasts, and mixotrophs, mirroring the trend in methanogen guild OTU richness (Figure 4, Figure S3). Hydrogenotrophs, acetoclasts, and methyl-reducers were generally positively associated with CH_4_ flux in all sites (Figure 5). These three guilds also tended to be positively correlated with salinity, while mixotrophic methanogens had mixed relationships among the sites (Figure S3). Of the 13 methanogenic families identified across the whole dataset, Methanobacteriaceae (containing hydrogenotrophs and methyl-reducers), was the most abundant, followed by Methanoregulaceae (hydrogenotrophs), Methanosaetaceae (acetoclasts), Methanomassiliicoccaceae (methyl-reducers), and Methanosarcinaceae (mixotrophs containing hydrogenotrophs, acetoclasts, methylotrophs, and methyl-reducers). Methanofastidiosaceae (hydrogenotrophs and methylotrophs (84)) increased in abundance with salinity in both field studies (Figure S2). The ratio of methanogens to methanotrophs (MG:MT) was positively correlated with CH_4_ flux in three sites, but only significantly in the higher sample size site (San Francisco) (Figure 5).

**Figure 5.**
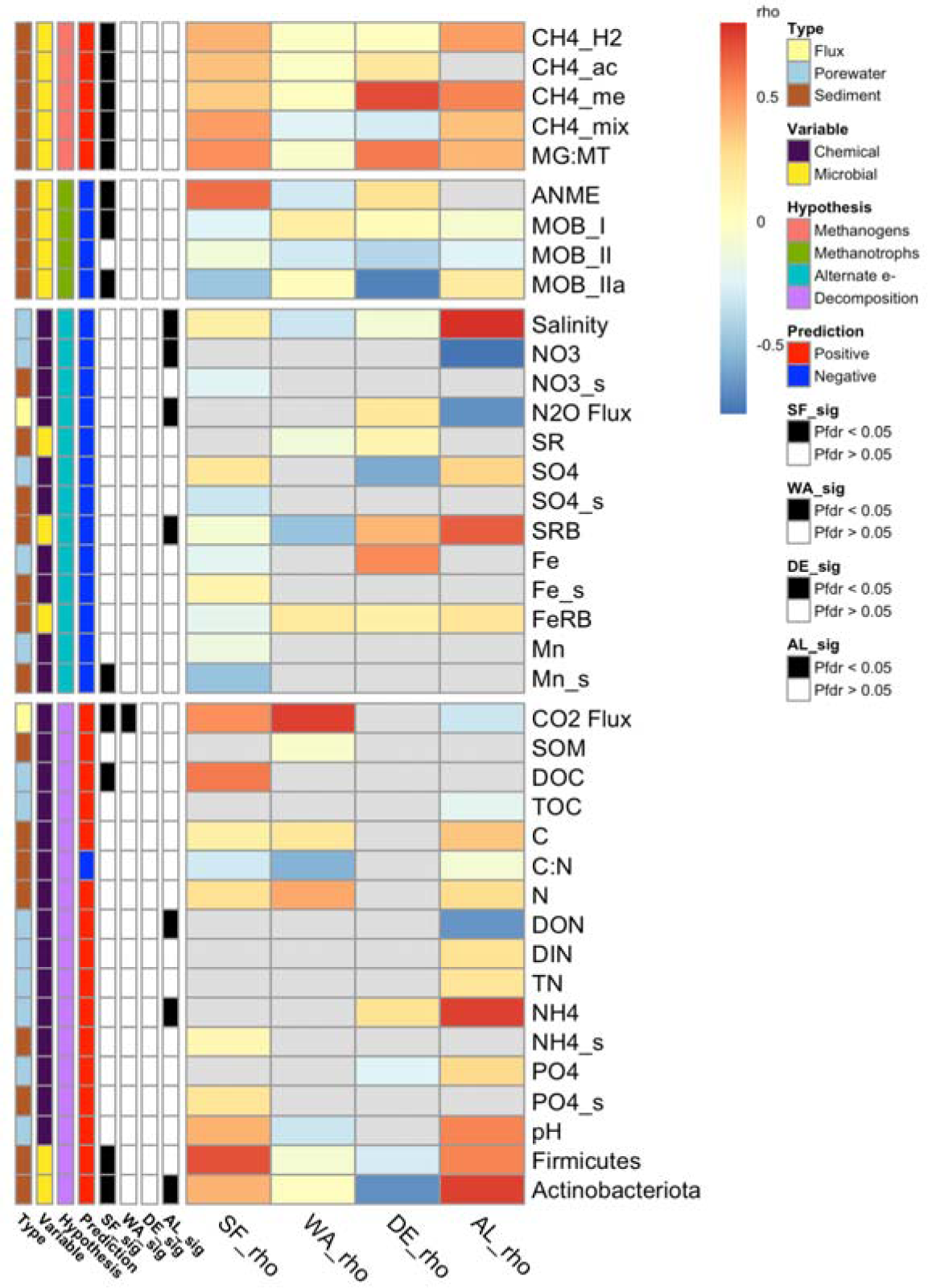
Correlations between chemical and microbial variables and methane flux. Rows are annotated by type (flux, porewater or soil), variable (chemical or microbial), hypothesis, the predicted relationship with methane (positive or negative), and whether the test was significant in each site (black) or not (white). The heatmap shows Spearman’s rho values and is broken into four panels, with the top two showing methanogenic and methanotrophic guilds and the bottom two corresponding to hypotheses about alternative electron acceptors and decomposition. Guild abbreviations are given in the Figure 3 caption. Gray cells indicate data not present. Data shown here from DE are from the field experiment only. MG:MT = methanogen: methanotroph ratio, SR = sulfate reduction rate; for chemical abbreviations see the methods text.

**Figure 6.**
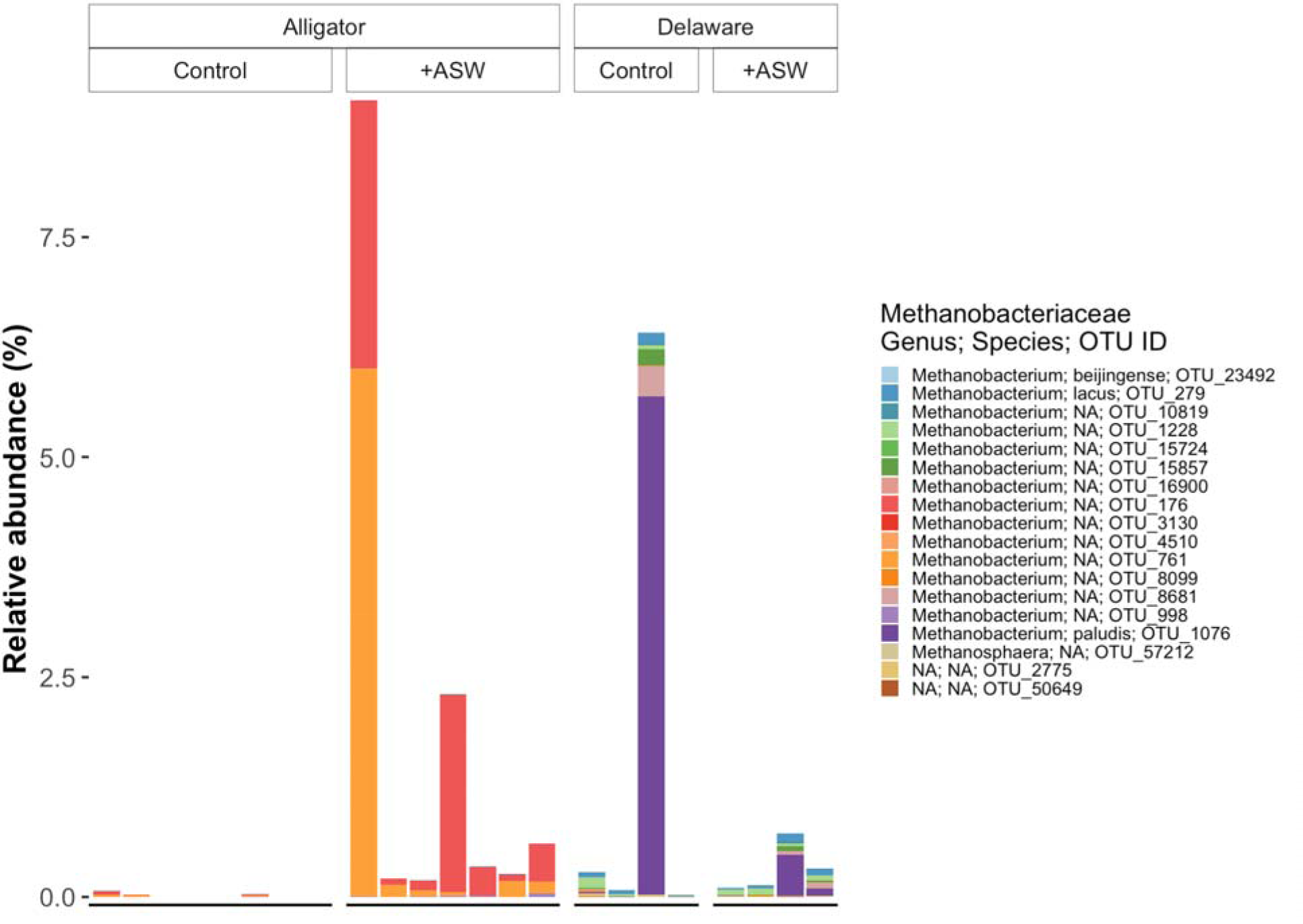
Methanobacteriaceae percent relative abundances for each individual sample in the laboratory incubation experiments conducted with wetland soils from the Alligator River and Delaware River estuaries.

There were 445 methanotroph OTUs: 394 MOB_I OTUs, 24 MOB_II OTUs, 21 MOB_IIa OTUs, and 6 ANME OTUs. Methanotroph community composition varied among the 4 sites (Figure S4). The aerobic class I methane oxidizing bacteria (MOB_I) were the most abundant methanotrophic guild across the whole dataset, followed by MOB_II, MOB_IIa, and ANME. The most abundant methanotrophic genera included *Methylocystis*, *Crenothrix*, *Methyloceanibacter*, and *Methylocaldum*. Some taxa consistently responded to salinity in the two field studies; in both San Francisco and Waccamaw, *Methyloceanibacter* increased with salinity while *Methylomagnum* and *Candidatus Methylospira* decreased with salinity (Figure S4). The four methanotrophic guilds had variable relationships with CH_4_ flux among the four sites (Figure 4). AO:NOB ratios were not significantly correlated with methanotroph abundances or and methanogen: methanotroph ratios except in San Francisco, where they were positively correlated, and Waccamaw, where the ratios were negatively correlated (Figure S5).

At the OTU level, there was a diverse group of 48 OTUs that could be considered strong indicators of either freshwater (n = 7) or oligohaline (n = 41) salinity conditions, with FDR corrected p-values < 0.05 and indicator correlation coefficients > 0.45 (Figure S6). Freshwater indicator taxa included OTUs from 7 different phyla but none were classified to genus. Oligohaline indicator OTUs included taxa from 13 different phyla and encompass some known genera and functional guild classifications such as the sulfate reducers *Desulfatiglans* and *SEEP-SRB1* and the nitrifiers *Nitrospira*, as well as taxa only identified to broader taxonomic levels.

Several biogeochemical variables and microbial guilds were significantly correlated with methane flux (Figure 5). In terms of salinity and alternative electron acceptors expected to be negatively correlated with CH_4_ flux, NO_3_^-^ and N_2_O flux were negatively correlated with CH_4_ flux in the Alligator incubation, and Mn was negatively correlated with CH_4_ flux in San Francisco. Salinity as a continuous variable (as opposed to categorical in Figure 1) was significantly *positively* correlated with CH_4_ flux in Alligator incubations and otherwise uncorrelated. For metrics of soil fertility and decomposition rates, CO_2_ flux, total soil C and N, and porewater NH_4_^+^ concentrations were generally positively correlated with CH_4_ flux, though not in all sites in which the variable was measured. pH was not significantly correlated with CH_4_ flux in any of the three sites in which it was measured. Several variables were significantly correlated with salinity, including a positive correlation between NH_4_^+^ and salinity in both Delaware and Alligator, but contrasting relationships between salinity and N_2_O flux at those two sites (Figure S2).

A key contrasting result was an increase in CH_4_ flux following ASW addition in the Alligator River estuary forested wetland laboratory incubations, and lack of increased CH_4_ flux in week 12 of the Delaware River estuary laboratory incubations (Figure 1), both of which were performed using similar methodology. Soils from Alligator have low pHs (< 5.5, Figure S7) and were dominated to a much greater extent by Acidobacteriota (Figure 4). In terms of the dominant methanogenic taxa, different OTUs within the Methanobacteriaceae family were dominant at each site, and these different OTUs responded differently to the +ASW treatment. Two OTUs at Alligator increased with ASW addition and were associated with greater CH_4_ fluxes (Figure 7).

The percent of shared OTUs among salinity classes within each site was 44-45% in laboratory experiments and 34-40% in field samples (Figure S8). Nearest taxon index (NTI) was > 2 in 131 of 133 samples (Figure S9). NTI was significantly lower in oligohaline samples than freshwater samples in San Francisco, Waccamaw, and Alligator but not Delaware (Figure S9). Dissimilarity between freshwater and oligohaline samples versus samples in the same salinity class was greater for both Bray-Curtis and Jaccard metrics; Cohen’s *d* effect size was slightly greater for Jaccard dissimilarity, but this was not affected by field or lab conditions (Figure S10).

## Discussion

### Microbial Alpha-Diversity (H1)

Salinity is expected to decrease the alpha diversity of freshwater microbial communities as increased salinity creates osmotic stress that requires organisms to either produce compatible solutes or, at extreme salinities, have specialized systems to function at high internal salt concentrations (19, 20). While there are many organisms that can tolerate a range of salinities, oligohaline salinities are high enough to cause stress or mortality in those organisms that have not evolved the ability to cope with a range of salinities (85). Our data partially support this hypothesis, as we found a negative effect of oligohaline salinity on richness in San Francisco and Waccamaw; however, richness in Delaware and Alligator was not affected. Decreases in bacterial richness have been found elsewhere, such as in freshwater streams affected by mining-induced salinity (86) and along salinity gradients in coastal wetlands (87). In contrast, slight increases in salinity can also cause an increase in richness, which has been found in Louisiana wetlands up to 3.5 ppt salinity (88), in Tibetan lakes up to 1 ppt salinity (89), and in Chinese soils exposed to up to 3.33 ppt irrigation water (90). Other work has suggested that bacterial communities are resistant to changes up to 3 ppt in salinity (91).

### Microbial Beta-Diversity and Taxa (H2)

Microbial community composition is strongly linked to enzyme activity and is thus important to understand with respect to biogeochemistry (92). Globally, salinity is a primary determinant of bacterial and archaeal community composition (16, 93), so it was not surprising to find differences between freshwater and oligohaline communities in all five studies here, although the effect of salinity was secondary to the effect of site (Figure 2). The broad differences in communities among the four sites are likely a product of different site histories, plant communities, and biogeochemistry (Table 1, Figure S7). Despite these differences, there were many (50 phyla, 121 classes, 237 orders, 264 families, 291 genera) shared taxa among the four sites (Figure S11). The indicator species analysis also suggested that there are some taxa shared across the sites that consistently associate with either freshwater or oligohaline salinities. A high proportion of Proteobacteria was found in all sites and is consistent with other studies in coastal wetlands (87, 88, 94). The decline in Verrucomicrobiota in oligohaline sites has also been observed across the Baltic Sea salinity gradient (95) and demonstrates a possible preference of this phylum for non-saline environments, although some members have been cultured at seawater salinities (96). Myxococcota have been identified as key components of tidal freshwater wetlands elsewhere (31), and have been shown to decline with increased salinity (97), in concordance with our results. While we found positive effects of salinity on the relative abundance of Chloroflexi, another common freshwater and estuarine wetland phylum (98, 99), results in other studies have shown mixed responses to low-level salinity (87, 100). Firmicutes, the other major phylum with consistent responses to salinity in more than one site, are also dominant members of other freshwater wetlands and have been shown to respond positively to salinity across estuarine (100), groundwater (101), and upland soil (102, 103) salinity gradients.

### Methanogens and Methanotrophs (H3)

While tidal wetlands have high functional redundancy for CO_2_ production, there are stronger links between CH_4_ production and microbes due to the conserved nature of methanogenic archaea (62). All four methanogenic guilds distinguishable with taxonomic data were present in the dataset, and the most abundant methanogens were hydrogenotrophs and acetoclasts, suggesting that these pathways of methanogenesis dominate in these wetlands. All guilds were generally positively associated with CH_4_ flux, except for mixotrophs in Waccamaw and Delaware. This discrepancy is likely explained by other controls on CH_4_ flux at play in the samples with high mixotroph abundance. In the Delaware River Estuary, isotope-based measurements demonstrated that acetoclastic methanogenesis dominated over hydrogenotrophic methanogenesis, although both contributed to the observed CH_4_ flux; methyl-based methanogenesis was not estimated (30, 69). Acetoclastic methanogens identified in the Delaware 16S data include the pure acetoclastic genus *Methanosaeta* and the mixotrophic genus *Methanosarcina*. On the other hand, pure acetoclasts were absent from Alligator; acetoclastic methanogenesis may still contribute to CH_4_ flux there via mixotrophic Methanosarcinaceae, which were present and positively correlated with CH_4_ flux. Acetoclasts have been suggested to increase with low salinity (104); in our datasets this was only the case in San Francisco. Though not as abundant, taxa in the methyl-reducing Massilliicococcales order were present in all sites and positively correlated with CH_4_ flux (though only significantly in San Francisco). The relative contribution of methyl-based methanogenesis is expected to increase with increasing salinity and in marine to hypersaline conditions due to the non-competitive substrates (e.g., betaine, trimethylamine) produced as osmolytes (50, 105, 106), but is perhaps less relevant at low salinities. In publicly available metagenomic datasets from freshwater and coastal wetlands, the abundance of archaeal genes involved in different methanogenesis pathways followed similar trends of a gene involved in acetoclastic methanogenesis (*cdhD*) being the most abundant, followed by hydrogenotrophic (*frhA*) and methylotrophic genes, with higher abundances in freshwater wetlands (50). The ratio of methanogens to methanotrophs (MG:MT) was positively correlated with CH_4_ flux in most sites, as seen elsewhere (107, 108), suggesting that this is an important microbial metric to calculate. Furthermore, MG:MT was positively correlated with salinity in Alligator soils, in line with the increase in CH_4_ flux there.

A substantial proportion of the total CH_4_ produced in methanogenic environments is consumed by methanotrophs (i.e., methane oxidizers) before it is emitted to the atmosphere, making them an important factor in regulating net CH_4_ fluxes. In marine soils the amount has been estimated to range from 43 - 85% (109). Four methanotrophic guilds were present at each site, yet their abundances were not consistently negatively correlated with CH_4_ flux, highlighting the complexity of factors that contribute to observed CH_4_ fluxes. MOB_I and MOB_IIa were negatively correlated with CH_4_ flux in San Francisco, however, and their higher abundances may have suppressed emissions in a wetland complex with higher methanogen abundances (23). Prior work has shown that both aerobic and anaerobic methanotrophy declined with salinity, with particular sensitivity of aerobic methanotrophy (110). In our dataset, aerobic bacterial methanotrophs (MOB_I, MOB_II, MOB_IIa) were much more abundant than anaerobic archaeal methanotrophs (ANME). Two of the three aerobic guilds did decline with salinity, but only in San Francisco (Figure 3), so our results only weakly agree with prior work (110). Methanotroph communities differed among the sites (Figure S4), with key abundant taxa including *Crenothrix* and *Methylocystis*. *Crenothrix* spp. are also dominant methanotrophs in stratified lakes (111), while *Methylocystis* spp. are also dominant in peatlands (112). While some *Methylocystis* species are inhibited by salinities > 5 ppt (113), *Methylocystis* was still abundant in Waccamaw oligohaline samples. *Methylospira* and *Methylomagnum* declined with salinity in both field studies, while *Methyloceanibacter*, containing known marine methanotrophs (114), not surprisingly increased with salinity, highlighting the taxa-specific environmental preferences of methanotrophs to different salinities. This may explain the lack of relationship between aggregated guild abundances and salinity, as at the guild level, all of the potentially positively and negatively responding taxa are summed.

Previous work in rice paddies has suggested links between nitrogen cyclers and methanotrophs (48, 57, 115). AOA and AOB perform the first step of nitrification, oxidizing NH_3_ to NO_2_^-^. Then NOB perform the second step, oxidizing NO_2_^-^ to NO_3_^-^. Thus, the combined activities of AO and NOB create NO_3_^-^, which could suppress CH_4_ flux due to the stimulation of nitrate reducers which can outcompete methanogens or by the inhibition of methane oxidizers. Alternatively, AO and NOB could increase CH_4_ flux due to increased inorganic N available for organic carbon-producing plant growth and microbial growth. Furthermore, a high ratio of AO to NOB could indicate nitrite buildup, which, along with other forms of inorganic N, could be toxic to methanotrophs in excess (59, 115). Our abundance data for AO:NOB and methanotrophs did not support this hypothesis of inhibition, with variation among sites and a lack of relationship in most cases. While influenced by agricultural runoff, our sites are unfertilized and therefore have lower inorganic nitrogen levels than the agricultural settings studied in previous work on inorganic N inhibition of methanotrophs (58).

While the results on methanogens and methanotrophs presented here are limited to 16S rRNA marker gene sequencing, they agree with shotgun metagenomic data from two of the sites – San Francisco and Alligator (23, 116). In particular, relative abundances of methanogens and methanotrophs from 16S rRNA gene PCR amplicons matched relative abundances of 16S rRNA genes extracted from metagenomes with mTAGs (116, 117), suggesting limited effects of any potential primer biases. Furthermore, relative abundances of *mcrA* and *amoA*, key marker genes of methanogenesis and methanotrophy, respectively, annotated from metagenomic data by sequence homology (118), were strongly and significantly correlated with the relative abundances of methanogens and methanotrophs extracted taxonomically from 16S PCR amplicon data. While we did not perform qPCR of these key genes, the homology-based approach with shotgun metagenomic data was actually more strongly correlated with 16S taxonomic relative abundances than *in silico* PCR, which was assessed for *mcrA* and *amoA* (23).

### CH_4_, Salinity, Alternative e-acceptors (H4)

While methane fluxes are generally expected to decrease with low salinity, as shown by several meta-analyses (37–39) and studies (68, 104, 119–121), data from a diverse range of individual sites and experiments show that salinity/methane relationships are highly variable and include positive, negative, and neutral relationships (41, 42). The degree of salinity and length of exposure to salinity are important, with different relationships seen above and below 7.5 ppt (122) and over different time periods; for example, previous work at Waccamaw demonstrated that long-term saltwater intrusion reduced both CO_2_ and CH_4_ flux, while short-term exposure increased CO_2_ but decreased CH_4_ (25). This is also an important source of variation among our 5 studies, where samples had been exposed to salinity for decades (San Francisco), 3 years (Waccamaw), 1 year (Delaware field), and 3 months (Delaware lab, Alligator lab) (Figure 5). Elsewhere, after 1 week of saltwater exposure CO_2_ increased but CH_4_ decreased (94). These results may be attributable to microbial community composition changes and adaptations. Intrusion events followed by recovery from salinity involve dynamic microbial interactions that affect carbon cycling (104). Temporary changes in salinity are expected to select for different microbial taxa (e.g., adapted to a range of salinities) compared to long-term changes in salinity, which are expected to select for taxa optimized for growth at the given salinity (120). For example, when freshwater and marine microbial communities were mixed, the community shifted towards the marine community composition, and when freshwater communities were exposed to sterile marine water, more freshwater taxa were lost from the community than saltwater taxa were lost when saltwater communities were exposed to freshwater (123). Another important variable is hydrologic setting – in permanently flooded soils, low salinity suppressed methanogenesis, while in intermittently flooded soils, low salinity increased methanogenesis (42). Flooded samples have greater abundances of both methanogens and methanotrophs than non-flooded samples (107). However, in our dataset all samples were continuously covered with water so that factor is not a source of variation here. We sought to use additional data on microbial community composition and biogeochemistry to explain these discrepancies.

Seawater exposure is expected to increase the availability of terminal electron acceptors for microbial growth, which affects the pathways by which organic matter is degraded (17, 124). In particular, nitrate, sulfate, iron, and manganese are key electron acceptors for microbial metabolism in anaerobic environments, and in tidal freshwater marshes in particular (17, 46, 125). We hypothesized that salinization by seawater would suppress CH_4_ flux due to competition between organisms reducing those electron acceptors and methanogens. Furthermore, these acceptors are used by methanotrophs during methane oxidation, whose activity would further decrease CH_4_ flux (55). Nitrate was negatively correlated with CH_4_ emissions at Alligator, consistent with what has been shown in rice paddies (58). This is also consistent with findings that increases in NO_3_^-^ increase denitrification rates (126), and denitrifier abundances increase with low salinity (127). Fluxes of N_2_O, a product of denitrification of nitrate, were significantly negatively correlated with CH_4_ flux at Alligator, consistent with potential competitive interactions between denitrifiers and methanogens. Of the alternative electron acceptors, sulfate has been the most widely implicated as a control on methanogenesis in freshwater (33) and estuarine (51) environments. However, here we find no support for suppression of CH_4_ production by sulfate at oligohaline salinities in the sites studied. SRB were the most abundant microbial functional guild and several sulfate reducing taxa were indicators of oligohaline conditions. However, SRB were actually *positively* correlated with CH_4_ flux in Alligator soils. Relationships between iron and manganese and salinity are less clear; both increased with salinity in our datasets, but this could also be caused by changes in oxidation states. However, there is no evidence in our data that increased Fe-reducing bacteria led to suppression of methanogenesis. Elsewhere, salinity has initially temporarily increased iron reduction rates, but this effect diminished over time (17). Another alternative electron acceptor in wetland soils is humic substances, which are used by certain anaerobic methanotrophs (ANME) (128), certain sulfate reducers (129), and other taxa (130). The abundance of humic substances could suppress CH_4_ emissions by either increasing activity of ANME (which consume CH_4_), increasing the activity of SRB and other humic substance reducers that can outcompete methanogens, or by causing methanogens to switch their metabolism from methanogenesis to humic substance reduction, which does not produce methane (131). We did not quantify humic and phenolic compounds in our studies, but we hypothesize they could be important in the forested and more acidic Alligator site, as SUVA_254_, a common metric of aromaticity and humics in water, was negatively correlated with CH_4_ flux earlier in the same experiment on which we report here (41); this remains an important avenue for future research.

While we only present results here from week 12 of the lab experiments, for which there was microbial data available, results from Alligator (increase in CH_4_ following artificial seawater addition) were consistent with the entire time course of the experiment (41). CH_4_ emissions were significantly elevated starting at day 21 of the experiment and continuing until the experiment’s conclusion on day 112. While using the limited amount of flux data available from cores with microbial data (week 12) from Delaware did not show elevated CH_4_ flux (Figure 1), the complete time series of that experiment showed increased CH_4_ emissions within one week after artificial seawater addition and continuing for 5 months (69).

### CH_4_ and Decomposition (H5)

There are multiple mechanisms by which low salinity can directly and indirectly affect decomposition rates in general, not just methanogenic decomposition. This can impact the directionality of salinity/C mineralization relationships, which have been shown to be highly variable in studies that have quantified either gas flux, enzymatic activities, or mass loss (40). For example, low salinity has had positive (17, 68, 69, 92, 119–121, 132–135) and negative (43, 136–141) effects on CO_2_ production, positive (92, 142) and negative (88) effects on enzymatic activities, and a negative effect on mass loss (143). Saltwater intrusion affects organic carbon production and solubility, which indirectly affect mineralization by controlling availability (25). Low salinity can decrease plant productivity (12) and C solubility (43), which would decrease overall mineralization rates. In addition to influencing carbon solubility, seawater can affect inorganic N (particularly ammonium) and P sorption, liberating nutrients, which in turn affect microbial activity and decomposition rates (26, 27, 144). Low salinity can either increase (145) or decrease (43, 122) pH, which similarly affects the solubility of compounds, the sorption of phosphorus, and decomposition rates. pH is also a strong driver of microbial community composition in both terrestrial (146) and aquatic (147) ecosystems. Lower pH is associated with, and can be driven by, a higher concentration of recalcitrant humics and phenolics that could slow decomposition rates and methanogenesis, as discussed above (131).

We predicted that regardless of how they were affected by salinity, CH_4_ fluxes would be coupled with CO_2_ flux and metrics of organic C, which should fuel metabolic pathways producing CO_2_ and CH_4_ (124, 148, 149). We also predicted positive correlations between CH_4_ flux and ammonium and phosphate concentrations, pH, and the relative abundance of Firmicutes and Actinobacteriota. Ammonium and phosphate are important nutrients for microbial growth, low pH may inhibit certain methanogens (150), and Firmicutes and Actinobacteriota are known to degrade complex carbon sources, which may supply substrates for methanogenesis (23, 151). There was mixed support for these predictions. CO_2_ and CH_4_ were indeed coupled in two field study sites (San Francisco, Waccamaw), but not in Alligator. Notably, although CO_2_ is generally produced in much higher quantities than CH_4_, it is also produced during acetoclastic and methylotrophic methanogenesis, but not hydrogenotrophic methanogenesis and the most abundant methanogens at Alligator were hydrogenotrophs. Different C, N, and phosphorus (P) variables were measured among the sites/experiments, but there were some instances of positive correlations with CH_4_ flux, such as DOC in San Francisco, and NH_4_^+^ in Alligator soils.

Ammonium can either increase or decrease CH_4_ emissions based on the balance between its effects at the plant/ecosystem level, the microbial community level, and the biochemical level (152). In Alligator soils, it was associated with increased CH_4_ flux, possibly due to increased N available for plant and methanogen growth (41). Phosphate was measured in two sites but was not significantly correlated with CH_4_ flux, suggesting that it does not exert a strong influence on CH_4_ in these coastal wetlands. Notably, these sites are likely not phosphate limited, which may partially explain this lack of effect; in P limited systems phosphate abundance may be more important (153, 154). Methanogenesis relies on upstream depolymerization of polymeric organic matter, as well as degradation of fatty acids such as butyrate (104). In this way methanogenesis is coupled to both photosynthesis and complex carbon degrading microorganisms. Metagenomic data has demonstrated positive correlations between hemicellulose and cellulose degrading genes and CH_4_ flux, and that Firmicutes and Actinobacteriota were the dominant taxonomic groups containing those genes (23). Firmicutes and Actinobacteriota were associated with increased CH_4_ flux at both San Francisco and Alligator. They have also been implicated in the depolymerization of rice straw, particularly in cellulose and hemicellulose degradation, fueling methanogenesis in rice paddies (151, 155); Actinobacteriota were also associated with methane production in *Sphagnum* peat bogs (156). A lack of relationship in Delaware and Waccamaw could be due to the lower overall abundance of Firmicutes in these sites; other taxa, including those in the Proteobacteria and Bacteroidetes phyla, may be responsible for complex C degradation in those locations.

### Site-Specificity

What could be driving such discrepancies in CH_4_/salinity relationships among the sites/experiments? One clear difference is the low pH and high abundance of Acidobacteriota and Firmicutes in Alligator laboratory-incubated soils compared to Delaware laboratory-incubated soils (Figure 3, Figure S7). This is likely driven by broad differences in plant community composition between the sites, which affects organic carbon quality and quantity which in turn affect microbial communities and decomposition rates (124, 125, 157, 158).

However, another laboratory experiment with soils from a tidal forested wetland similar to Alligator showed a decrease in CH_4_ production with seawater addition, highlighting the need for more studies in forested wetlands to be able to make generalizations (121). Alligator also notably had the smallest and least rich methanogen population in all samples, particularly in oligohaline samples (Figure S4). While humic substances were not quantified, they are expected to be higher in the acidic Alligator wetland compared to Delaware Estuary wetlands, and this could suggest another mechanism for increased CH_4_ at that site in particular.

There is also support for the idea that site-specific microbial consortia that are functionally relevant have different responses to low salinity. While many broader taxonomic groups were shared among the four sites, relatively few were shared at the OTU level (Figure S11). A case in point are the methanogens in the most abundant methanogen family, Methanobacteriaceae (Figure 7). Most Methanobacteriaceae reads were from the hydrogenotrophic *Methanobacterium* genus, but even within the genus there were different OTUs and some evidence for different responses to oligohaline salinity. The idea that different species in the same genus, or even different strains of the same species, have different environmental preferences is not new, especially when considering pathogens (159), and has been greatly expanded upon in recent years with the help of full genome sequencing and metabolic modeling. For example, metabolic modeling of 55 *E. coli* strains showed differences in niches among the strains (160). Such site-specific OTU niche differences and salinity responses, particularly of taxa involved in CH_4_ cycling, can then contribute to discrepancies in CH_4_/salinity relationships.

### Field vs. Lab Settings

Another potential source of discrepancy in CH_4_/salinity relationships among the five datasets examined here could arise from differences in field and laboratory experimental conditions. Field samples are, not surprisingly, characterized by greater richness and diversity than the laboratory samples taken after three months in the laboratory (Figure 1). Laboratory experiments also did not include plants and therefore do not take into account changes in C inputs as plants grow, release exudates, and senesce. To test for differences in the ecological dynamics between laboratory and field samples, we analyzed taxa overlap and nearest taxon index (NTI), and compared a presence/absence metric (Jaccard) with an abundance-based metric (Bray-Curtis). Field samples are expected to have more unique taxa in freshwater and oligohaline conditions than samples incubated in the lab; field samples are mixed with the surrounding environment and microbes can disperse into the sampled soil from the surrounding environment, whereas samples incubated in the laboratory are cut off from the regional species pool after they are initially collected from the field. Dispersal and immigration are key processes in microbial community assembly (161). Residence times of estuarine microbes in the field are on the order of days to weeks, indicating the potential for dynamic temporal turnover (162).

As expected, laboratory experiments had a higher degree of overlap in OTUs among salinity classes (Figure S8). However, differences in community composition among salinity classes were similar when using presence/absence and abundance-based metrics, and there were no differences among field and lab samples (Figure S10). Furthermore, virtually all samples, whether from field or lab, were dominated by deterministic assembly processes (Figure S9), even in the field where stochasticity could have potentially played a larger role due to dispersal and immigration. The NTI values > 2 suggest phylogenetic clustering in most samples. Overall this suggests that similar ecological dynamics are at play in the lab in the field, namely the deterministic processes of selection, competition, and fitness differences among species (163). However, the difference in alpha diversity is notable and may affect the degree of microbial community functional redundancy, and consequently the effects of salinity on C mineralization.

## Conclusions

Salinization is a major issue caused by sea level rise, storms and tides, drought, water management, and water-body connectivity, which leads to increased ionic strength, alkalinization, and sulfidation, which can result in coastal forest loss, species invasion, yield declines, eutrophication, marsh migration, and changes in microbial communities and biogeochemical cycling (13). An outstanding question regarding salinization of coastal wetlands is whether CH_4_ emissions will increase or decrease, with important implications for C storage and climate change (41). Our synthesis of microbial and biogeochemical data from multiple studies and methods from multiple geographic locations highlights both some consistencies in the wetland soil microbial communities that regulate CH_4_ cycling, as well as some important differences that can qualitatively affect how CH_4_ emissions respond to increases in low salinity, including microbial community responses and length of exposure. Importantly, our results do not support the assumption that seawater intrusion will generally decrease CH_4_ emissions from coastal wetlands, at least in the short-term. Further systematic paired sampling of microbial communities and CH_4_ flux measurements across a greater number of sites exposed to the same amount of salinity for the same amount of time is needed to directly analyze CH_4_/salinity relationship as a response variable, and to be able to make more generalizations about CH_4_/salinity relationships in tidal freshwater marshes and other coastal and estuarine wetlands.

## Acknowledgements

We acknowledge the JGI sequencing staff for processing the samples and performing the DNA sequencing. We thank all of the team members who helped collect samples in the field and helped run the laboratory incubation experiments. The work conducted by the U.S. Department of Energy Joint Genome Institute, a DOE Office of Science User Facility, is supported by the Office of Science of the U.S. Department of Energy under Contract No. DE-AC02-05CH11231.

## Open Research

Raw data and analysis scripts for this submission are publicly available on Zenodo (https://doi.org/10.5281/zenodo.8250416) (164). Sequencing data are available on NCBI GenBank with BioProject ID PRJNA1004999 (165).

## References

1. IPCC. 2021. Climate Change 2021: The Physical Science Basis. Contribution of Working Group I to the Sixth Assessment Report of the Intergovernmental Panel on Climate Change. Cambridge University Press.

2. Etheridge DM, Steele LP, Francey RJ, Langenfelds RL. 1998. Atmospheric methane between 1000 A.D. and present: Evidence of anthropogenic emissions and climatic variability. J Geophys Res Atmospheres 103:15979–15993.

3. Saunois M, Stavert AR, Poulter B, Bousquet P, Canadell JG, Jackson RB, Raymond PA, Dlugokencky EJ, Houweling S, Patra PK, Ciais P, Arora VK, Bastviken D, Bergamaschi P, Blake DR, Brailsford G, Bruhwiler L, Carlson KM, Carrol M, Castaldi S, Chandra N, Crevoisier C, Crill PM, Covey K, Curry CL, Etiope G, Frankenberg C, Gedney N, Hegglin MI, Höglund-Isaksson L, Hugelius G, Ishizawa M, Ito A, Janssens-Maenhout G, Jensen KM, Joos F, Kleinen T, Krummel PB, Langenfelds RL, Laruelle GG, Liu L, Machida T, Maksyutov S, McDonald KC, McNorton J, Miller PA, Melton JR, Morino I, Müller J, Murguia-Flores F, Naik V, Niwa Y, Noce S, O’Doherty S, Parker RJ, Peng C, Peng S, Peters GP, Prigent C, Prinn R, Ramonet M, Regnier P, Riley WJ, Rosentreter JA, Segers A, Simpson IJ, Shi H, Smith SJ, Steele LP, Thornton BF, Tian H, Tohjima Y, Tubiello FN, Tsuruta A, Viovy N, Voulgarakis A, Weber TS, van Weele M, van der Werf GR, Weiss RF, Worthy D, Wunch D, Yin Y, Yoshida Y, Zhang W, Zhang Z, Zhao Y, Zheng B, Zhu Q, Zhu Q, Zhuang Q. 2020. The global methane budget 2000–2017. Earth Syst Sci Data 12:1561–1623.

4. Nellemann C. 2009. Blue carbon. A UNEP rapid response assessment.

5. Grimsditch G, Alder J, Nakamura T, Kenchington R, Tamelander J. 2013. The blue carbon special edition – Introduction and overview. Ocean Coast Manag 83:1–4.

6. Mcleod E, Chmura GL, Bouillon S, Salm R, Björk M, Duarte CM, Lovelock CE, Schlesinger WH, Silliman BR. 2011. A blueprint for blue carbon: toward an improved understanding of the role of vegetated coastal habitats in sequestering CO2. Front Ecol Environ 9:552–560.

7. Kelleway JJ, Serrano O, Baldock JA, Burgess R, Cannard T, Lavery PS, Lovelock CE, Macreadie PI, Masqué P, Newnham M, Saintilan N, Steven ADL. 2020. A national approach to greenhouse gas abatement through blue carbon management. Glob Environ Change 63:102083.

8. Valach AC, Kasak K, Hemes KS, Szutu D, Verfaillie J, Baldocchi DD. 2021. Carbon flux trajectories and site conditions from restored impounded marshes in the Sacramento-San Joaquin Delta, p. 247–271. In Wetland Carbon and Environmental Management. First Edition. John Wiley & Sons, Inc.

9. Hemes KS, Chamberlain SD, Eichelmann E, Knox SH, Baldocchi DD. 2018. A Biogeochemical Compromise: The High Methane Cost of Sequestering Carbon in Restored Wetlands. Geophys Res Lett 45:6081–6091.

10. Rosentreter JA, Al-Haj AN, Fulweiler RW, Williamson P. 2021. Methane and Nitrous Oxide Emissions Complicate Coastal Blue Carbon Assessments. Glob Biogeochem Cycles 35:e2020GB006858.

11. Herbert ER, Boon P, Burgin AJ, Neubauer SC, Franklin RB, Ardón M, Hopfensperger KN, Lamers LPM, Gell P. 2015. A global perspective on wetland salinization: ecological consequences of a growing threat to freshwater wetlands. Ecosphere 6:art206.

12. Chamberlain SD, Hemes KS, Eichelmann E, Szutu DJ, Verfaillie JG, Baldocchi DD. 2020. Effect of Drought-Induced Salinization on Wetland Methane Emissions, Gross Ecosystem Productivity, and Their Interactions. Ecosystems 23:675–688.

13. Tully K, Gedan K, Epanchin-Niell R, Strong A, Bernhardt ES, BenDor T, Mitchell M, Kominoski J, Jordan TE, Neubauer SC, Weston NB. 2019. The Invisible Flood: The Chemistry, Ecology, and Social Implications of Coastal Saltwater Intrusion. BioScience 69:368–378.

14. Crain CM, Silliman BR, Bertness SL, Bertness MD. 2004. Physical and Biotic Drivers of Plant Distribution Across Estuarine Salinity Gradients. Ecology 85:2539–2549.

15. Zervoudaki S, Nielsen TG, Carstensen J. 2009. Seasonal succession and composition of the zooplankton community along an eutrophication and salinity gradient exemplified by Danish waters. J Plankton Res 31:1475–1492.

16. Lozupone CA, Knight R. 2007. Global patterns in bacterial diversity. Proc Natl Acad Sci 104:11436–11440.

17. Weston NB, Dixon RE, Joye SB. 2006. Ramifications of increased salinity in tidal freshwater sediments: Geochemistry and microbial pathways of organic matter mineralization. J Geophys Res Biogeosciences 111.

18. Mohamed DJ, Martiny JB. 2011. Patterns of fungal diversity and composition along a salinity gradient. 3. ISME J 5:379–388.

19. Gunde-Cimerman N, Plemenitaš A, Oren A. 2018. Strategies of adaptation of microorganisms of the three domains of life to high salt concentrations. FEMS Microbiol Rev 42:353–375.

20. Oren A. 2013. Life at high salt concentrations, intracellular KCl concentrations, and acidic proteomes. Front Microbiol 0.

21. Odum WE. 1988. Comparative Ecology of Tidal Freshwater and Salt Marshes. Annu Rev Ecol Syst 19:147–176.

22. Dragone NB, Diaz MA, Hogg ID, Lyons WB, Jackson WA, Wall DH, Adams BJ, Fierer N. 2021. Exploring the Boundaries of Microbial Habitability in Soil. J Geophys Res Biogeosciences 126:e2020JG006052.

23. Hartman WH, Bueno de Mesquita CP, Theroux SM, Morgan-Lang C, Baldocchi DD, Tringe SG. 2024. Multiple microbial guilds mediate soil methane cycling along a wetland salinity gradient. mSystems e00936–23.

24. Laas P, Ugarelli K, Travieso R, Stumpf S, Gaiser EE, Kominoski JS, Stingl U. 2022. Water Column Microbial Communities Vary along Salinity Gradients in the Florida Coastal Everglades Wetlands. 2. Microorganisms 10:215.

25. Neubauer SC, Franklin RB, Berrier DJ. 2013. Saltwater intrusion into tidal freshwater marshes alters the biogeochemical processing of organic carbon. Biogeosciences 10:8171–8183.

26. Ardón M, Morse JL, Colman BP, Bernhardt ES. 2013. Drought-induced saltwater incursion leads to increased wetland nitrogen export. Glob Change Biol 19:2976–2985.

27. Weston NB, Giblin AE, Banta GT, Hopkinson CS, Tucker J. 2010. The Effects of Varying Salinity on Ammonium Exchange in Estuarine Sediments of the Parker River, Massachusetts. Estuaries Coasts 33:985–1003.

28. Hu M, Peñuelas J, Sardans J, Yang X, Tong C, Zou S, Cao W. 2020. Shifts in Microbial Biomass C/N/P Stoichiometry and Bacterial Community Composition in Subtropical Estuarine Tidal Marshes Along a Gradient of Freshwater–Oligohaline Water. Ecosystems 23:1265–1280.

29. Lew S, Glińska-Lewczuk K, Burandt P, Kulesza K, Kobus S, Obolewski K. 2022. Salinity as a Determinant Structuring Microbial Communities in Coastal Lakes. 8. Int J Environ Res Public Health 19:4592.

30. Weston NB, Neubauer SC, Velinsky DJ, Vile MA. 2014. Net ecosystem carbon exchange and the greenhouse gas balance of tidal marshes along an estuarine salinity gradient. Biogeochemistry 120:163–189.

31. Morina JC, Franklin RB. 2022. Intensity and duration of exposure determine prokaryotic community response to salinization in freshwater wetland soils. Geoderma 428:116138.

32. Achtnich C, Bak F, Conrad R. 1995. Competition for electron donors among nitrate reducers, ferric iron reducers, sulfate reducers, and methanogens in anoxic paddy soil. Biol Fertil Soils 19:65–72.

33. Lovley DR, Klug MJ. 1983. Sulfate Reducers Can Outcompete Methanogens at Freshwater Sulfate Concentrations. Appl Environ Microbiol 45:187–192.

34. Kristjansson JK, Schönheit P. 1983. Why do sulfate-reducing bacteria outcompete methanogenic bacteria for substrates? Oecologia 60:264–266.

35. Lovley DR, Dwyer DF, Klug MJ. 1982. Kinetic Analysis of Competition Between Sulfate Reducers and Methanogens for Hydrogen in Sediments. Appl Environ Microbiol 43:1373– 1379.

36. Schönheit P, Kristjansson JK, Thauer RK. 1982. Kinetic mechanism for the ability of sulfate reducers to out-compete methanogens for acetate. Arch Microbiol 132:285–288.

37. Bartlett KB, Bartlett DS, Harriss RC, Sebacher DI. 1987. Methane emissions along a salt marsh salinity gradient. Biogeochemistry 4:183–202.

38. Poffenbarger HJ, Needelman BA, Megonigal JP. 2011. Salinity influence on methane emissions from tidal marshes. Wetlands 31:831–842.

39. Al-Haj AN, Fulweiler RW. 2020. A synthesis of methane emissions from shallow vegetated coastal ecosystems. Glob Change Biol 26:2988–3005.

40. Luo M, Huang J-F, Zhu W-F, Tong C. 2019. Impacts of increasing salinity and inundation on rates and pathways of organic carbon mineralization in tidal wetlands: a review. Hydrobiologia 827:31–49.

41. Ardón M, Helton AM, Bernhardt ES. 2018. Salinity effects on greenhouse gas emissions from wetland soils are contingent upon hydrologic setting: a microcosm experiment. Biogeochemistry 140:217–232.

42. Helton AM, Ardón M, Bernhardt ES. 2019. Hydrologic Context Alters Greenhouse Gas Feedbacks of Coastal Wetland Salinization. Ecosystems 22:1108–1125.

43. Ury EA, Wright JP, Ardón M, Bernhardt ES. 2022. Saltwater intrusion in context: soil factors regulate impacts of salinity on soil carbon cycling. Biogeochemistry 157:215–226.

44. Ardón M, Helton AM, Scheuerell MD, Bernhardt ES. 2017. Fertilizer legacies meet saltwater incursion: challenges and constraints for coastal plain wetland restoration. Elem Sci Anthr 5.

45. Hopple AM, Pennington SC, Megonigal JP, Bailey V, Bond-Lamberty B. 2022. Disturbance legacies regulate coastal forest soil stability to changing salinity and inundation: A soil transplant experiment. Soil Biol Biochem 169:108675.

46. Lovley DR. 1991. Dissimilatory Fe(III) and Mn(IV) reduction. Microbiol Rev 55:259–287.

47. Conrad R. 2020. Importance of hydrogenotrophic, aceticlastic and methylotrophic methanogenesis for methane production in terrestrial, aquatic and other anoxic environments: A mini review. Pedosphere 30:25–39.

48. Bodelier PL, Steenbergh AK. 2014. Interactions between methane and the nitrogen cycle in light of climate change. Curr Opin Environ Sustain 9–10:26–36.

49. Kurth JM, Op den Camp HJM, Welte CU. 2020. Several ways one goal—methanogenesis from unconventional substrates. Appl Microbiol Biotechnol 104:6839–6854.

50. Bueno de Mesquita CP, Wu D, Tringe SG. 2023. Methyl-Based Methanogenesis: an Ecological and Genomic Review. Microbiol Mol Biol Rev 0:e00024–22.

51. Oremland RS, Polcin S. 1982. Methanogenesis and sulfate reduction: competitive and noncompetitive substrates in estuarine sediments. Appl Environ Microbiol 44:1270–1276.

52. Oremland RS, Marsh LM, Polcin S. 1982. Methane production and simultaneous sulphate reduction in anoxic, salt marsh sediments. 5853. Nature 296:143–145.

53. Xu L, Zhuang G-C, Montgomery A, Liang Q, Joye SB, Wang F. 2021. Methyl-compounds driven benthic carbon cycling in the sulfate-reducing sediments of South China Sea. Environ Microbiol 23:641–651.

54. Maltby J, Sommer S, Dale AW, Treude T. 2016. Microbial methanogenesis in the sulfate-reducing zone of surface sediments traversing the Peruvian margin. Biogeosciences 13:283–299.

55. Guerrero-Cruz S, Vaksmaa A, Horn MA, Niemann H, Pijuan M, Ho A. 2021. Methanotrophs: Discoveries, Environmental Relevance, and a Perspective on Current and Future Applications. Front Microbiol 12.

56. Conrad R. 2007. Microbial Ecology of Methanogens and Methanotrophs, p. 1–63. In Advances in Agronomy. Academic Press.

57. Bodelier PL. 2011. Interactions between nitrogenous fertilizers and methane cycling in wetland and upland soils. Curr Opin Environ Sustain 3:379–388.

58. Cai Z, Shan Y, Xu H. 2007. Effects of nitrogen fertilization on CH4 emissions from rice fields. Soil Sci Plant Nutr 53:353–361.

59. Dunfield P, Knowles R. 1995. Kinetics of inhibition of methane oxidation by nitrate, nitrite, and ammonium in a humisol. Appl Environ Microbiol 61:3129–3135.

60. Alam M, Jia Z. 2012. Inhibition of methane oxidation by nitrogenous fertilizers in a paddy soil. Front Microbiol 3:246.

61. Nyerges G, Stein LY. 2009. Ammonia cometabolism and product inhibition vary considerably among species of methanotrophic bacteria. FEMS Microbiol Lett 297:131– 136.

62. Morrissey EM, Berrier DJ, Neubauer SC, Franklin RB. 2014. Using microbial communities and extracellular enzymes to link soil organic matter characteristics to greenhouse gas production in a tidal freshwater wetland. Biogeochemistry 117:473–490.

63. Luo Z, Wang E, Smith C. 2015. Fresh carbon input differentially impacts soil carbon decomposition across natural and managed systems. Ecology 96:2806–2813.

64. Yan W, Zhong Y, Zhu G, Liu W, Shangguan Z. 2020. Nutrient limitation of litter decomposition with long-term secondary succession: evidence from controlled laboratory experiments. J Soils Sediments 20:1858–1868.

65. Barantal S, Schimann H, Fromin N, Hättenschwiler S. 2012. Nutrient and Carbon Limitation on Decomposition in an Amazonian Moist Forest. Ecosystems 15:1039–1052.

66. Walse C, Berg B, Sverdrup H. 1998. Review and synthesis of experimental data on organic matter decomposition with respect to the effect of temperature, moisture, and acidity. Environ Rev 6:25–40.

67. Ardón M, Morse JL, Doyle MW, Bernhardt ES. 2010. The Water Quality Consequences of Restoring Wetland Hydrology to a Large Agricultural Watershed in the Southeastern Coastal Plain. Ecosystems 13:1060–1078.

68. Neubauer SC. 2013. Ecosystem Responses of a Tidal Freshwater Marsh Experiencing Saltwater Intrusion and Altered Hydrology. Estuaries Coasts 36:491–507.

69. Weston NB, Vile MA, Neubauer SC, Velinsky DJ. 2011. Accelerated microbial organic matter mineralization following salt-water intrusion into tidal freshwater marsh soils. Biogeochemistry 102:135–151.

70. Tremblay J, Singh K, Fern A, Kirton ES, He S, Woyke T, Lee J, Chen F, Dangl JL, Tringe SG. 2015. Primer and platform effects on 16S rRNA tag sequencing. Front Microbiol 6.

71. Callahan BJ, McMurdie PJ, Rosen MJ, Han AW, Johnson AJA, Holmes SP. 2016. DADA2: High-resolution sample inference from Illumina amplicon data. Nat Methods 13:581–583.

72. Quast C, Pruesse E, Yilmaz P, Gerken J, Schweer T, Yarza P, Peplies J, Glöckner FO. 2013. The SILVA ribosomal RNA gene database project: improved data processing and web-based tools. Nucleic Acids Res 41:D590–D596.

73. Leff JW. 2017. mctoolsr: Microbial Community Data Analysis Tools (0.1.1.2). R.

74. Oksanen J, Blanchet FG, Friendly M, Kindt R, Legendre P, McGlinn D, Minchin PR, O’Hara RB, Simpson GL, Solymos P, Stevens MHH, Szoecs E, Wagner H. 2019. vegan: Community Ecology Package. R package version 2.5–6. https://CRAN.R-project.org/package=vegan. (2.5-6) R.

75. De Cáceres M, Legendre P. 2009. Associations between species and groups of sites: indices and statistical inference. Ecology 90:3566–3574.

76. Edgar RC. 2010. Search and clustering orders of magnitude faster than BLAST. Bioinformatics 26:2460–2461.

77. Price MN, Dehal PS, Arkin AP. 2009. FastTree: Computing Large Minimum Evolution Trees with Profiles instead of a Distance Matrix. Mol Biol Evol 26:1641–1650.

78. Caporaso JG, Kuczynski J, Stombaugh J, Bittinger K, Bushman FD, Costello EK, Fierer N, Peña AG, Goodrich JK, Gordon JI, Huttley GA, Kelley ST, Knights D, Koenig JE, Ley RE, Lozupone CA, McDonald D, Muegge BD, Pirrung M, Reeder J, Sevinsky JR, Turnbaugh PJ, Walters WA, Widmann J, Yatsunenko T, Zaneveld J, Knight R. 2010. QIIME allows analysis of high-throughput community sequencing data. 5. Nat Methods 7:335–336.

79. Ning D, Yuan M, Wu L, Zhang Y, Guo X, Zhou X, Yang Y, Arkin AP, Firestone MK, Zhou J. 2020. A quantitative framework reveals ecological drivers of grassland microbial community assembly in response to warming. 1. Nat Commun 11:4717.

80. Stegen JC, Lin X, Konopka AE, Fredrickson JK. 2012. Stochastic and deterministic assembly processes in subsurface microbial communities. 9. ISME J 6:1653–1664.

81. Wickham H. 2016. ggplot2: Elegant Graphics for Data Analysis. Springer-Verlag, New York, NY. https://ggplot2.tidyverse.org.

82. Kolde R. 2019. pheatmap: Pretty Heatmaps. R package version 1.0.12. https://CRAN.R-project.org/package=pheatmap. (R package version 1.0.12) R.

83. R Core Team. 2023. R: A Language and Environment for Statistical Computing (4.2.3). R. R Foundation for Statistical Computing, Vienna, Austria.

84. Nobu MK, Narihiro T, Kuroda K, Mei R, Liu W-T. 2016. Chasing the elusive Euryarchaeota class WSA2: genomes reveal a uniquely fastidious methyl-reducing methanogen. ISME J 10:2478–2487.

85. Georges des Aulnois M, Roux P, Caruana A, Réveillon D, Briand E, Hervé F, Savar V, Bormans M, Amzil Z. 2019. Physiological and Metabolic Responses of Freshwater and Brackish-Water Strains of Microcystis aeruginosa Acclimated to a Salinity Gradient: Insight into Salt Tolerance. Appl Environ Microbiol 85:e01614–19.

86. Vander Vorste R, Timpano AJ, Cappellin C, Badgley BD, Zipper CE, Schoenholtz SH. 2019. Microbial and macroinvertebrate communities, but not leaf decomposition, change along a mining-induced salinity gradient. Freshw Biol 64:671–684.

87. Zhao Q, Bai J, Gao Y, Zhao H, Zhang G, Cui B. 2020. Shifts in the soil bacterial community along a salinity gradient in the Yellow River Delta. Land Degrad Dev 31:2255– 2267.

88. Jackson CR, Vallaire SC. 2009. Effects of salinity and nutrients on microbial assemblages in Louisiana wetland sediments. Wetlands 29:277–287.

89. Wang J, Yang D, Zhang Y, Shen J, Gast C van der, Hahn MW, Wu Q. 2011. Do Patterns of Bacterial Diversity along Salinity Gradients Differ from Those Observed for Macroorganisms? PLOS ONE 6:e27597.

90. Chen L, Li C, Feng Q, Wei Y, Zheng H, Zhao Y, Feng Y, Li H. 2017. Shifts in soil microbial metabolic activities and community structures along a salinity gradient of irrigation water in a typical arid region of China. Sci Total Environ 598:64–70.

91. Berga M, Zha Y, Székely AJ, Langenheder S. 2017. Functional and Compositional Stability of Bacterial Metacommunities in Response to Salinity Changes. Front Microbiol 8.

92. Morrissey EM, Gillespie JL, Morina JC, Franklin RB. 2014. Salinity affects microbial activity and soil organic matter content in tidal wetlands. Glob Change Biol 20:1351–1362.

93. Auguet J-C, Barberan A, Casamayor EO. 2010. Global ecological patterns in uncultured Archaea. 2. ISME J 4:182–190.

94. Dang C, Morrissey EM, Neubauer SC, Franklin RB. 2019. Novel microbial community composition and carbon biogeochemistry emerge over time following saltwater intrusion in wetlands. Glob Change Biol 25:549–561.

95. Herlemann DP, Labrenz M, Jürgens K, Bertilsson S, Waniek JJ, Andersson AF. 2011. Transitions in bacterial communities along the 2000 km salinity gradient of the Baltic Sea. 10. ISME J 5:1571–1579.

96. Schlesner H, Jenkins C, Staley JT. 2006. The Phylum Verrucomicrobia: A Phylogenetically Heterogeneous Bacterial Group, p. 881–896. In Dworkin, M, Falkow, S, Rosenberg, E, Schleifer, K-H, Stackebrandt, E (eds.), The Prokaryotes: Volume 7: Proteobacteria: Delta, Epsilon Subclass. Springer, New York, NY.

97. Zhao X, Meng T, Jin S, Ren K, Cai Z, Cai B, Li S. 2023. The Salinity Survival Strategy of Chenopodium quinoa: Investigating Microbial Community Shifts and Nitrogen Cycling in Saline Soils. 12. Microorganisms 11:2829.

98. Ikenaga M, Guevara R, Dean AL, Pisani C, Boyer JN. 2010. Changes in Community Structure of Sediment Bacteria Along the Florida Coastal Everglades Marsh–Mangrove– Seagrass Salinity Gradient. Microb Ecol 59:284–295.

99. Huang J, Zhu J, Liu S, Luo Y, Zhao R, Guo F, Li B. 2022. Estuarine salinity gradient governs sedimentary bacterial community but not antibiotic resistance gene profile. Sci Total Environ 806:151390.

100. Zhang G, Bai J, Tebbe CC, Zhao Q, Jia J, Wang W, Wang X, Yu L. 2021. Salinity controls soil microbial community structure and function in coastal estuarine wetlands. Environ Microbiol 23:1020–1037.

101. Sang S, Zhang X, Dai H, Hu BX, Ou H, Sun L. 2018. Diversity and predictive metabolic pathways of the prokaryotic microbial community along a groundwater salinity gradient of the Pearl River Delta, China. 1. Sci Rep 8:17317.

102. Van Horn DJ, Okie JG, Buelow HN, Gooseff MN, Barrett JE, Takacs-Vesbach CD. 2014. Soil Microbial Responses to Increased Moisture and Organic Resources along a Salinity Gradient in a Polar Desert. Appl Environ Microbiol 80:3034–3043.

103. Zheng W, Xue D, Li X, Deng Y, Rui J, Feng K, Wang Z. 2017. The responses and adaptations of microbial communities to salinity in farmland soils: A molecular ecological network analysis. Appl Soil Ecol 120:239–246.

104. Berrier DJ, Neubauer SC, Franklin RB. 2022. Cooperative microbial interactions mediate community biogeochemical responses to saltwater intrusion in wetland soils. FEMS Microbiol Ecol 98:fiac019.

105. Oren A. 1999. Bioenergetic Aspects of Halophilism. Microbiol Mol Biol Rev 63:334–348.

106. Zhou J, Theroux SM, Bueno de Mesquita CP, Hartman WH, Tringe SG. 2021. Microbial drivers of methane emissions from unrestored industrial salt ponds. ISME J 16:284–295.

107. Rey-Sanchez C, Bohrer G, Slater J, Li Y-F, Grau-Andrés R, Hao Y, Rich VI, Davies GM. 2019. The ratio of methanogens to methanotrophs and water-level dynamics drive methane transfer velocity in a temperate kettle-hole peat bog. Biogeosciences 16:3207– 3231.

108. Zhang Y, Cui M, Duan J, Zhuang X, Zhuang G, Ma A. 2019. Abundance, rather than composition, of methane-cycling microbes mainly affects methane emissions from different vegetation soils in the Zoige alpine wetland. MicrobiologyOpen 8:e00699.

109. Reeburgh WS. 2007. Oceanic Methane Biogeochemistry. Chem Rev 107:486–513.

110. Dalal RC, Allen DE, Livesley SJ, Richards G. 2008. Magnitude and biophysical regulators of methane emission and consumption in the Australian agricultural, forest, and submerged landscapes: a review. Plant Soil 309:43–76.

111. Oswald K, Graf JS, Littmann S, Tienken D, Brand A, Wehrli B, Albertsen M, Daims H, Wagner M, Kuypers MM, Schubert CJ, Milucka J. 2017. Crenothrix are major methane consumers in stratified lakes. 9. ISME J 11:2124–2140.

112. Chen Y, Dumont MG, Neufeld JD, Bodrossy L, Stralis-Pavese N, McNamara NP, Ostle N, Briones MJI, Murrell JC. 2008. Revealing the uncultivated majority: combining DNA stable-isotope probing, multiple displacement amplification and metagenomic analyses of uncultivated Methylocystis in acidic peatlands. Environ Microbiol 10:2609–2622.

113. Dedysh SN, Belova SE, Bodelier PLE, Smirnova KV, Khmelenina VN, Chidthaisong A, Trotsenko YA, Liesack W, Dunfield PF. 2007. Methylocystis heyeri sp. nov., a novel type II methanotrophic bacterium possessing ‘signature’ fatty acids of type I methanotrophs. Int J Syst Evol Microbiol 57:472–479.

114. Takeuchi M, Katayama T, Yamagishi T, Hanada S, Tamaki H, Kamagata Y, Oshima K, Hattori M, Marumo K, Nedachi M, Maeda H, Suwa Y, Sakata S. 2014. Methyloceanibacter caenitepidi gen. nov., sp. nov., a facultatively methylotrophic bacterium isolated from marine sediments near a hydrothermal vent. Int J Syst Evol Microbiol 64:462–468.

115. Bodelier PLE, Laanbroek HJ. 2004. Nitrogen as a regulatory factor of methane oxidation in soils and sediments. FEMS Microbiol Ecol 47:265–277.

116. Bueno de Mesquita CP, Hartman WH, Ardón M, Tringe SG. In review. Disentangling the effects of sulfate and other seawater ions on microbial communities and greenhouse gas emissions in a coastal forested wetland. ISME Commun.

117. Salazar G, Ruscheweyh H-J, Hildebrand F, Acinas SG, Sunagawa S. 2021. mTAGs: taxonomic profiling using degenerate consensus reference sequences of ribosomal RNA genes. Bioinformatics 38:270–272.

118. Morgan-Lang C, McLaughlin R, Armstrong Z, Zhang G, Chan K, Hallam SJ. 2020. TreeSAPP: the Tree-based Sensitive and Accurate Phylogenetic Profiler. Bioinformatics 36:4706–4713.

119. Chambers LG, Reddy KR, Osborne TZ. 2011. Short-Term Response of Carbon Cycling to Salinity Pulses in a Freshwater Wetland. Soil Sci Soc Am J 75:2000–2007.

120. Chambers LG, Osborne TZ, Reddy KR. 2013. Effect of salinity-altering pulsing events on soil organic carbon loss along an intertidal wetland gradient: a laboratory experiment. Biogeochemistry 115:363–383.

121. Marton JM, Herbert ER, Craft CB. 2012. Effects of Salinity on Denitrification and Greenhouse Gas Production from Laboratory-incubated Tidal Forest Soils. Wetlands 32:347–357.

122. Wang C, Tong C, Chambers LG, Liu X. 2017. Identifying the Salinity Thresholds that Impact Greenhouse Gas Production in Subtropical Tidal Freshwater Marsh Soils. Wetlands 37:559–571.

123. Rocca JD, Simonin M, Bernhardt ES, Washburne AD, Wright JP. 2020. Rare microbial taxa emerge when communities collide: freshwater and marine microbiome responses to experimental mixing. Ecology 101:e02956.

124. Sutton-Grier ArianaE, Keller JK, Koch R, Gilmour C, Megonigal JP. 2011. Electron donors and acceptors influence anaerobic soil organic matter mineralization in tidal marshes. Soil Biol Biochem 43:1576–1583.

125. Megonigal JP, Neubauer SC. 2019. Chapter 19 - Biogeochemistry of Tidal Freshwater Wetlands, p. 641–683. In Perillo, GME, Wolanski, E, Cahoon, DR, Hopkinson, CS (eds.), Coastal Wetlands (Second Edition). Elsevier.

126. Morrissey EM, Franklin RB. 2015. Resource effects on denitrification are mediated by community composition in tidal freshwater wetlands soils. Environ Microbiol 17:1520– 1532.

127. Franklin RB, Morrissey EM, Morina JC. 2017. Changes in abundance and community structure of nitrate-reducing bacteria along a salinity gradient in tidal wetlands. Pedobiologia 60:21–26.

128. Bai Y-N, Wang X-N, Wu J, Lu Y-Z, Fu L, Zhang F, Lau T-C, Zeng RJ. 2019. Humic substances as electron acceptors for anaerobic oxidation of methane driven by ANME-2d. Water Res 164:114935.

129. Cervantes FJ, Bok FAM de, Duong-Dac T, Stams AJM, Lettinga G, Field JA. 2002. Reduction of humic substances by halorespiring, sulphate-reducing and methanogenic microorganisms. Environ Microbiol 4:51–57.

130. Coates JD, Ellis DJ, Blunt-Harris EL, Gaw CV, Roden EE, Lovley DR. 1998. Recovery of Humic-Reducing Bacteria from a Diversity of Environments. Appl Environ Microbiol 64:1504–1509.

131. Valenzuela EI, Cervantes FJ. 2021. The role of humic substances in mitigating greenhouse gases emissions: Current knowledge and research gaps. Sci Total Environ 750:141677.

132. Saviozzi A, Cardelli R, Di Puccio R. 2011. Impact of Salinity on Soil Biological Activities: A Laboratory Experiment. Commun Soil Sci Plant Anal 42:358–367.

133. Liu X, Ruecker A, Song B, Xing J, Conner WH, Chow AT. 2017. Effects of salinity and wet–dry treatments on C and N dynamics in coastal-forested wetland soils: Implications of sea level rise. Soil Biol Biochem 112:56–67.

134. Baldwin DS, Rees GN, Mitchell AM, Watson G, Williams J. 2006. The short-term effects of salinization on anaerobic nutrient cycling and microbial community structure in sediment from a freshwater wetland. Wetlands 26:455–464.

135. Chambers LG, Davis SE, Troxler T, Boyer JN, Downey-Wall A, Scinto LJ. 2014. Biogeochemical effects of simulated sea level rise on carbon loss in an Everglades mangrove peat soil. Hydrobiologia 726:195–211.

136. Kelley CA, Martens CS, Chanton JP. 1990. Variations in sedimentary carbon remineralization rates in the White Oak River estuary, North Carolina. Limnol Oceanogr 35:372–383.

137. Nyman JA, DeLaune RD. 1991. CO2emission and soil Eh responses to different hydrological conditions in fresh, brackish, and saline marsh soils. Limnol Oceanogr 36:1406–1414.

138. Smith CJ, DeLaune RD, Patrick WH. 1983. Carbon dioxide emission and carbon accumulation in coastal wetlands. Estuar Coast Shelf Sci 17:21–29.

139. DeLaune RD, Smith CJ, Patrick WH. 1983. Methane release from Gulf coast wetlands. Tellus B Chem Phys Meteorol 35:8–15.

140. Wilson BJ, Mortazavi B, Kiene RP. 2015. Spatial and temporal variability in carbon dioxide and methane exchange at three coastal marshes along a salinity gradient in a northern Gulf of Mexico estuary. Biogeochemistry 123:329–347.

141. Krauss KW, Whitbeck JL. 2012. Soil Greenhouse Gas Fluxes during Wetland Forest Retreat along the Lower Savannah River, Georgia (USA). Wetlands 32:73–81.

142. Cunha MA, Almeida MA, Alcântara F. 2000. Patterns of ectoenzymatic and heterotrophic bacterial activities along a salinity gradient in a shallow tidal estuary. Mar Ecol Prog Ser 204:1–12.

143. Roache MC, Bailey PC, Boon PI. 2006. Effects of salinity on the decay of the freshwater macrophyte, Triglochin procerum. Aquat Bot 84:45–52.

144. Zhou M, Butterbach-Bahl K, Vereecken H, Brüggemann N. 2017. A meta-analysis of soil salinization effects on nitrogen pools, cycles and fluxes in coastal ecosystems. Glob Change Biol 23:1338–1352.

145. Portnoy JW, Giblin AE. 1997. Biogeochemical Effects of Seawater Restoration to Diked Salt Marshes. Ecol Appl 7:1054–1063.

146. Lauber CL, Hamady M, Knight R, Fierer N. 2009. Pyrosequencing-Based Assessment of Soil pH as a Predictor of Soil Bacterial Community Structure at the Continental Scale. Appl Environ Microbiol 75:5111–5120.

147. Sadeghi J, Chaganti SR, Shahraki AH, Heath DD. 2021. Microbial community and abiotic effects on aquatic bacterial communities in north temperate lakes. Sci Total Environ 781:146771.

148. Seo J, Jang I, Gebauer G, Kang H. 2014. Abundance of Methanogens, Methanotrophic Bacteria, and Denitrifiers in Rice Paddy Soils. Wetlands 34:213–223.

149. Oikawa PY, Jenerette GD, Knox SH, Sturtevant C, Verfaillie J, Dronova I, Poindexter CM, Eichelmann E, Baldocchi DD. 2017. Evaluation of a hierarchy of models reveals importance of substrate limitation for predicting carbon dioxide and methane exchange in restored wetlands. J Geophys Res Biogeosciences 122:145–167.

150. Van Kessel JAS, Russell JB. 1996. The effect of pH on ruminal methanogenesis. FEMS Microbiol Ecol 20:205–210.

151. Gavande PV, Basak A, Sen S, Lepcha K, Murmu N, Rai V, Mazumdar D, Saha SP, Das V, Ghosh S. 2021. Functional characterization of thermotolerant microbial consortium for lignocellulolytic enzymes with central role of Firmicutes in rice straw depolymerization. 1. Sci Rep 11:3032.

152. Schimel J. 2000. Rice, microbes and methane. 6768. Nature 403:375–377.

153. Herbert ER, Schubauer-Berigan JP, Craft CB. 2020. Effects of 10 yr of nitrogen and phosphorus fertilization on carbon and nutrient cycling in a tidal freshwater marsh. Limnol Oceanogr 65:1669–1687.

154. Ket WA, Schubauer-Berigan JP, Craft CB. 2011. Effects of five years of nitrogen and phosphorus additions on a Zizaniopsis miliacea tidal freshwater marsh. Aquat Bot 95:17– 23.

155. Kausar H, Sariah M, Mohd Saud H, Zahangir Alam M, Razi Ismail M. 2011. Isolation and screening of potential actinobacteria for rapid composting of rice straw. Biodegradation 22:367–375.

156. Pankratov T, Dedysh S, Zavarzin G. 2006. The leading role of actinobacteria in aerobic cellulose degradation in Sphagnum peat bogs. Dokl Biol Sci Proc Acad Sci USSR Biol Sci Sect Transl Russ 410:428–30.

157. Shahbaz M, Kuzyakov Y, Sanaullah M, Heitkamp F, Zelenev V, Kumar A, Blagodatskaya E. 2017. Microbial decomposition of soil organic matter is mediated by quality and quantity of crop residues: mechanisms and thresholds. Biol Fertil Soils 53:287–301.

158. Bueno de Mesquita CP, Schmidt SK, Suding KN. 2019. Litter-driven feedbacks influence plant colonization of a high elevation early successional ecosystem. Plant Soil 444:71–85.

159. Thelaus J, Lundmark E, Lindgren P, Sjödin A, Forsman M. 2018. Galleria mellonella Reveals Niche Differences Between Highly Pathogenic and Closely Related Strains of Francisella spp. Front Cell Infect Microbiol 8.

160. Monk JM, Charusanti P, Aziz RK, Lerman JA, Premyodhin N, Orth JD, Feist AM, Palsson BØ. 2013. Genome-scale metabolic reconstructions of multiple Escherichia coli strains highlight strain-specific adaptations to nutritional environments. Proc Natl Acad Sci 110:20338–20343.

161. Sloan WT, Lunn M, Woodcock S, Head IM, Nee S, Curtis TP. 2006. Quantifying the roles of immigration and chance in shaping prokaryote community structure. Environ Microbiol 8:732–740.

162. Crump BC, Hopkinson CS, Sogin ML, Hobbie JE. 2004. Microbial Biogeography along an Estuarine Salinity Gradient: Combined Influences of Bacterial Growth and Residence Time. Appl Environ Microbiol 70:1494–1505.

163. Vellend M. 2010. Conceptual Synthesis in Community Ecology. Q Rev Biol 85:183–206.

164. Bueno de Mesquita CP. 2024. JGR Resubmission (Version 2.0.0) [Dataset]. Zenodo. 10.5281/zenodo.8250416

165. Bueno de Mesquita CP. 2023. Microbial ecology and site characteristics underlie differences in salinity-methane relationships in coastal wetlands [Dataset]. PRJNA1004999. NCBI. https://www.ncbi.nlm.nih.gov/bioproject/PRJNA1004999/

