## Supplemental Figures for "Microbial ecology and site characteristics underlie differences in salinity-methane relationships in coastal wetlands"

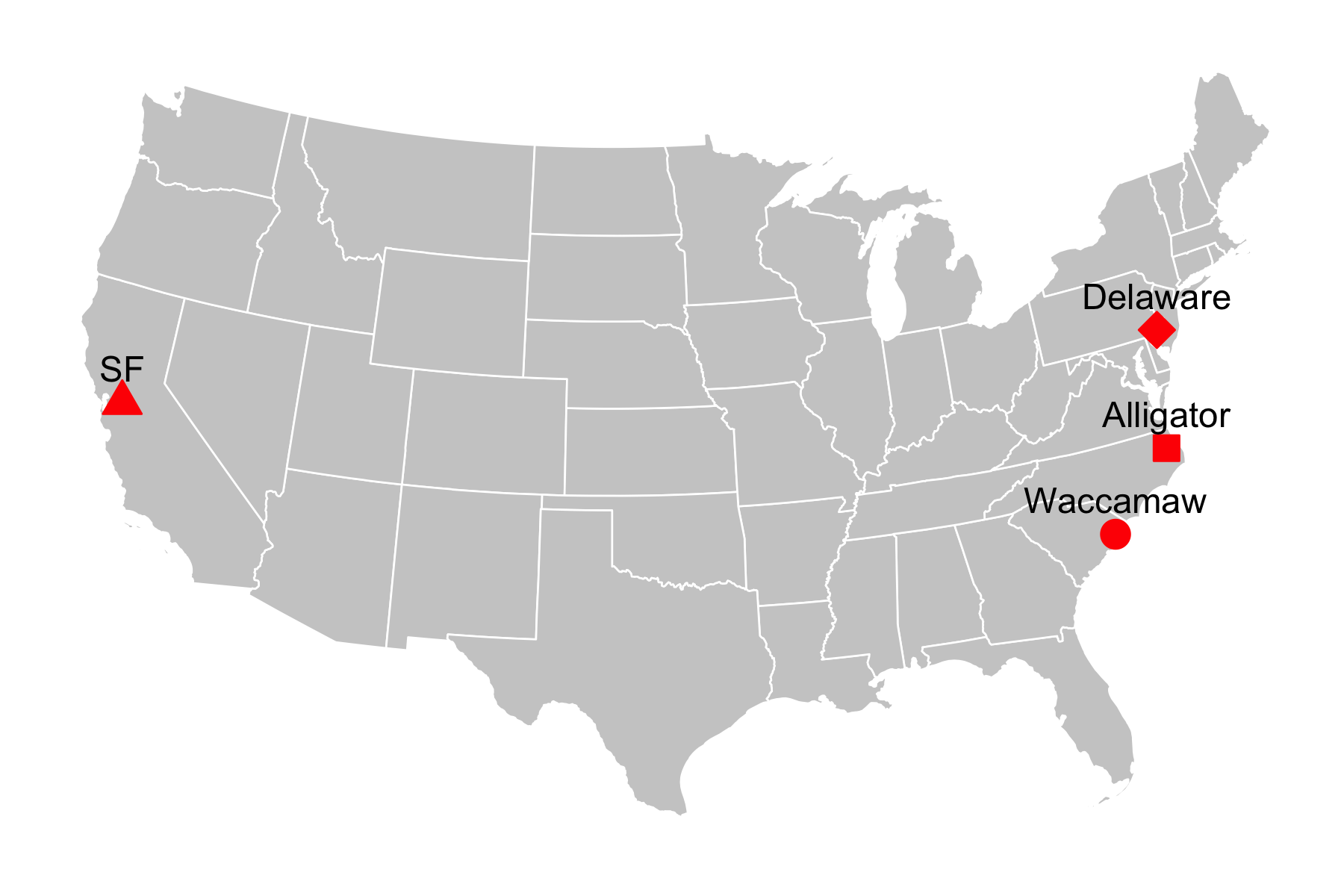

Figure S1. Map of the 4 broad study sites. SF and Delaware each contained 4 subsites (Table 1) while Alligator and Waccamaw each contained only 1 site.

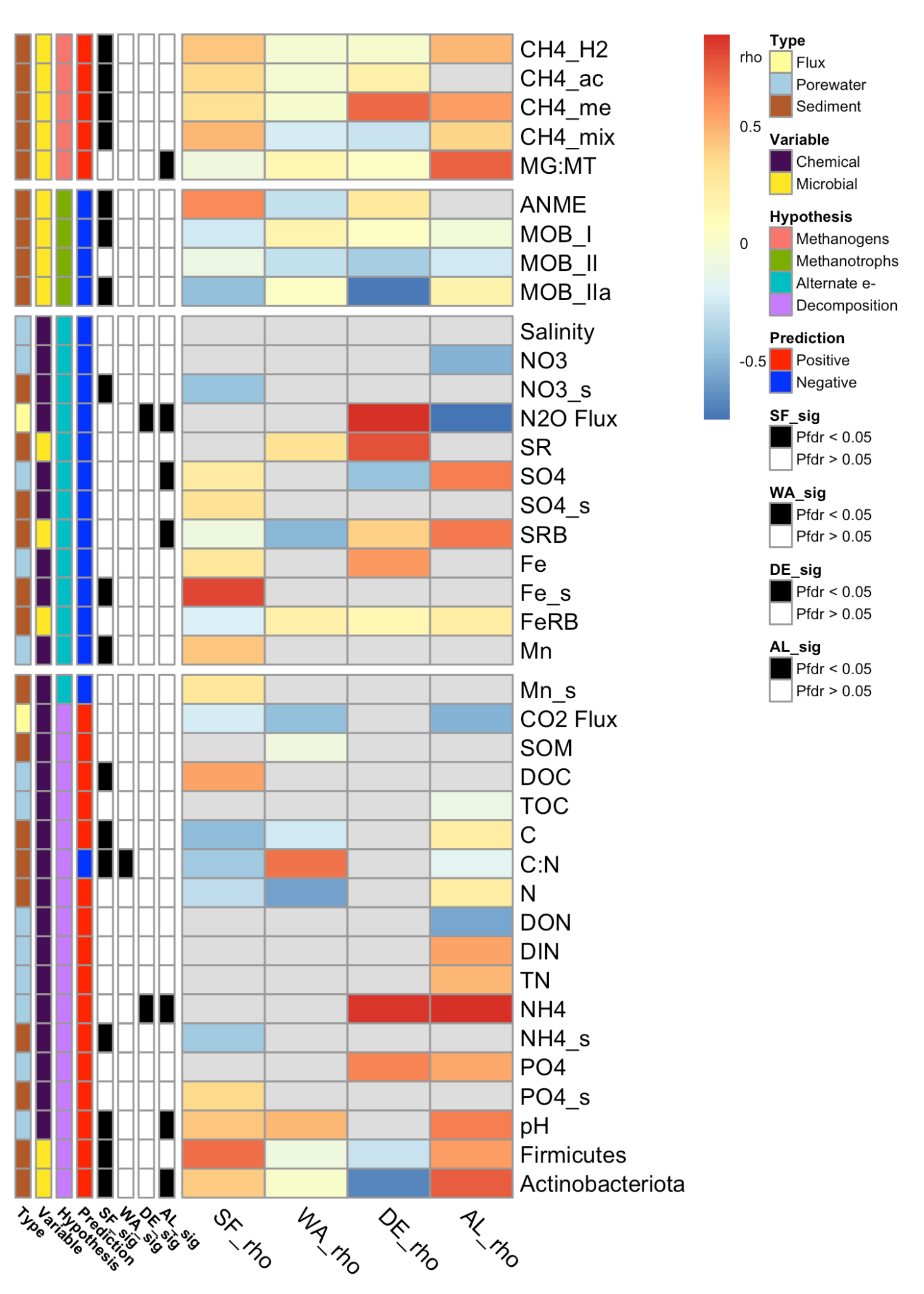

Figure S2. Correlations between chemical and microbial variables and salinity. Rows are annotated by type (flux, porewater or soil), variable (chemical or microbial), hypothesis, the predicted relationship with salinity (positive or negative), and whether the test was significant in each site (black) or not (white). The heatmap shows Spearman’s rho values and is broken into four panels, with the top two corresponding to hypotheses about alternative electron acceptors and decomposition, and the bottom two panels corresponding showing methanogenic and methanotrophic guilds. Guild abbreviations are given in the Figure 3 caption. Gray cells indicate data not present. Data shown here from DE are from the field experiment only. MG:MT = methanogen: methanotroph ratio, SR = sulfate reduction rate; for chemical abbreviations see the methods text.

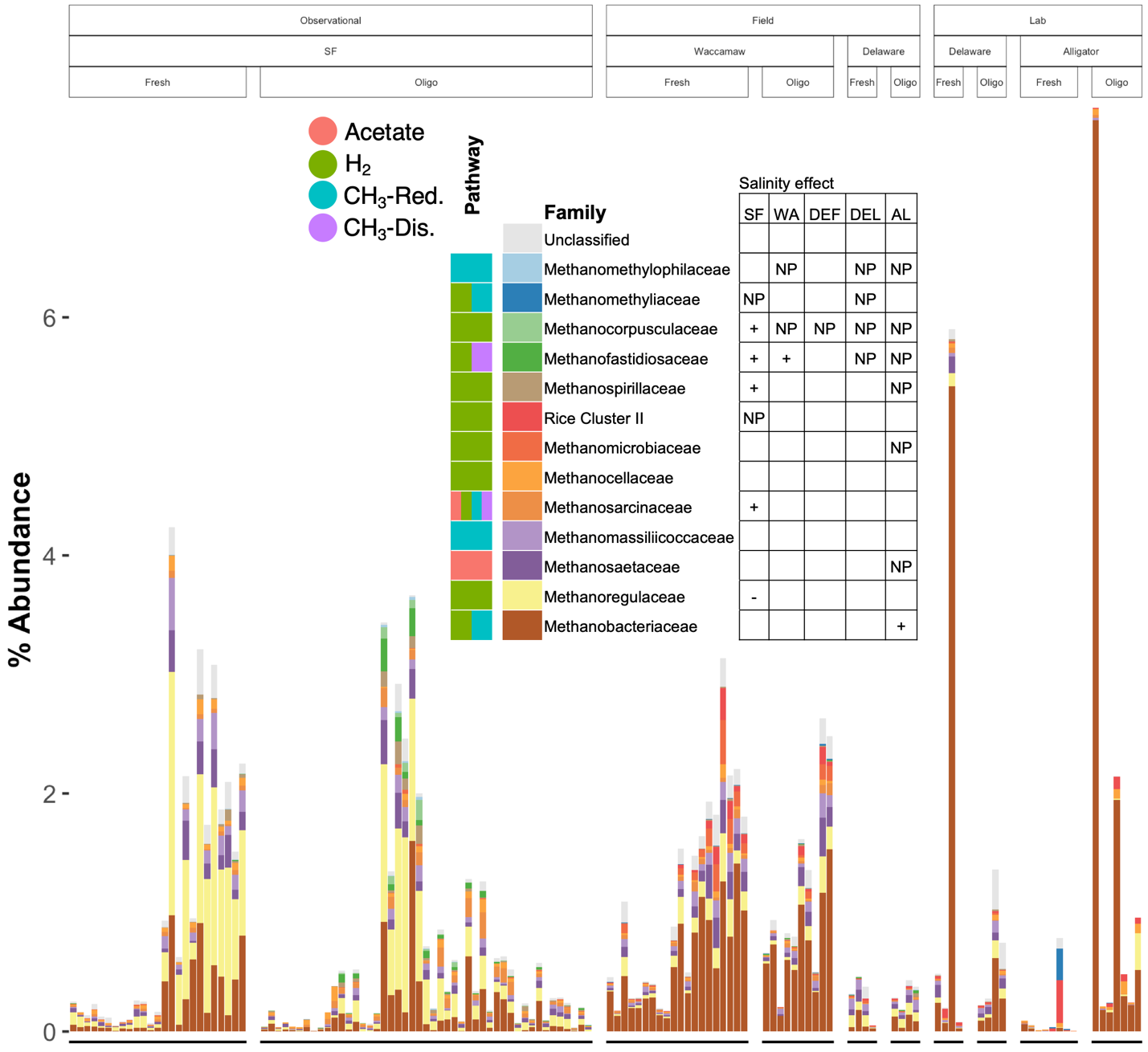
Figure S3. Relative abundances of methanogens, showing all 13 methanogenic families as well as the aggregated sum of all methanogenic taxa with taxonomy unclassified to the family level (“Unclassified”). Also shown is the methanogenic pathway(s) that members of the family are known to perform. NP = not present in a particular site/experiment; DEF = Delaware field experiment; DEL = Delaware lab experiment.

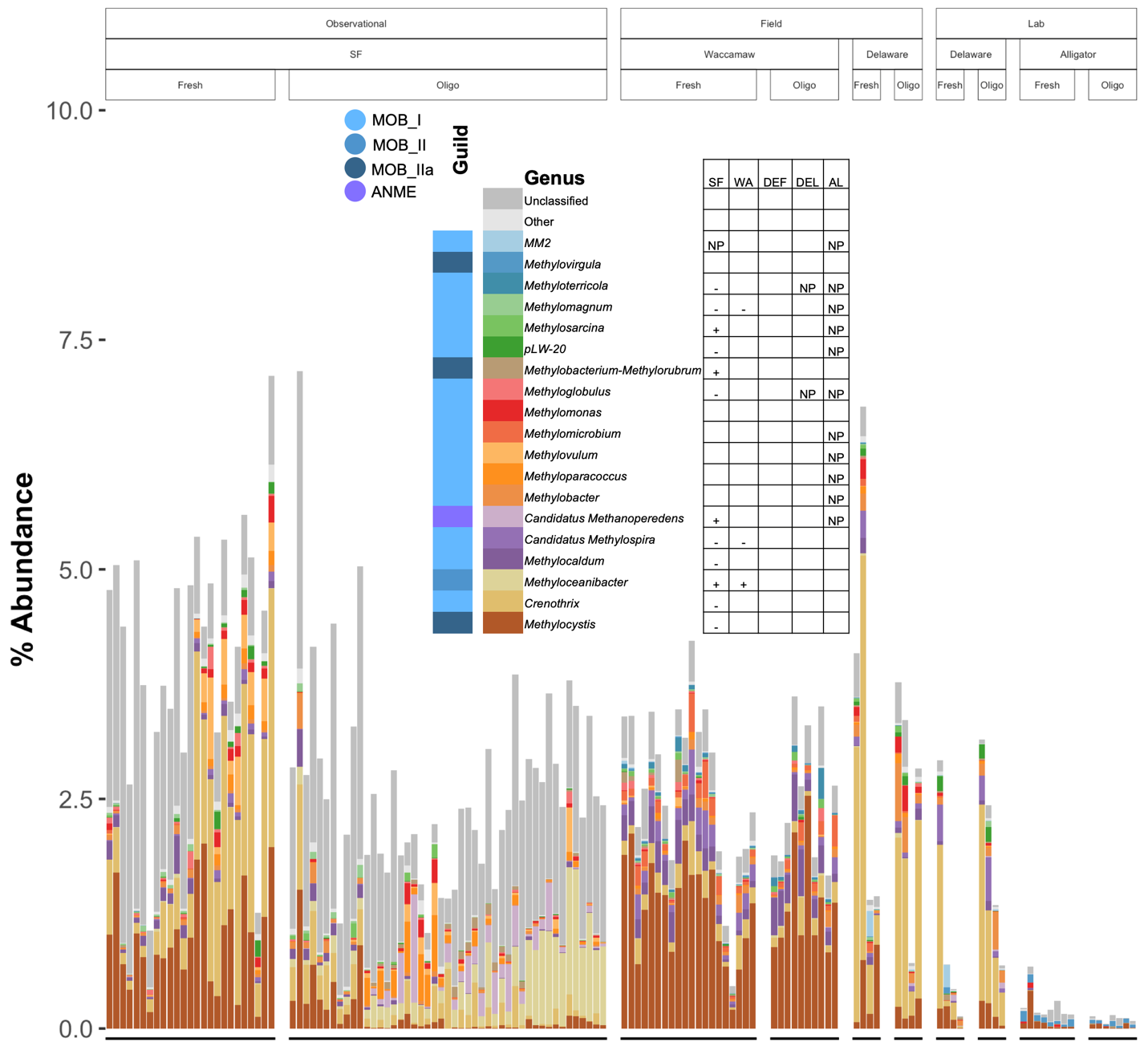
Figure S4. Relative abundance of methanotrophs (ANME + MOB_I + MOB_II + MOB_IIa), showing of top 19 most abundant methanotrophic genera, the aggregated sum of all other classified genera (“Other”), and the aggregated sum of all methanotrophic taxa unclassified to the genus level (“Unclassified”). NP = not present in a particular site/experiment; DEF = Delaware field experiment; DEL = Delaware lab experiment.

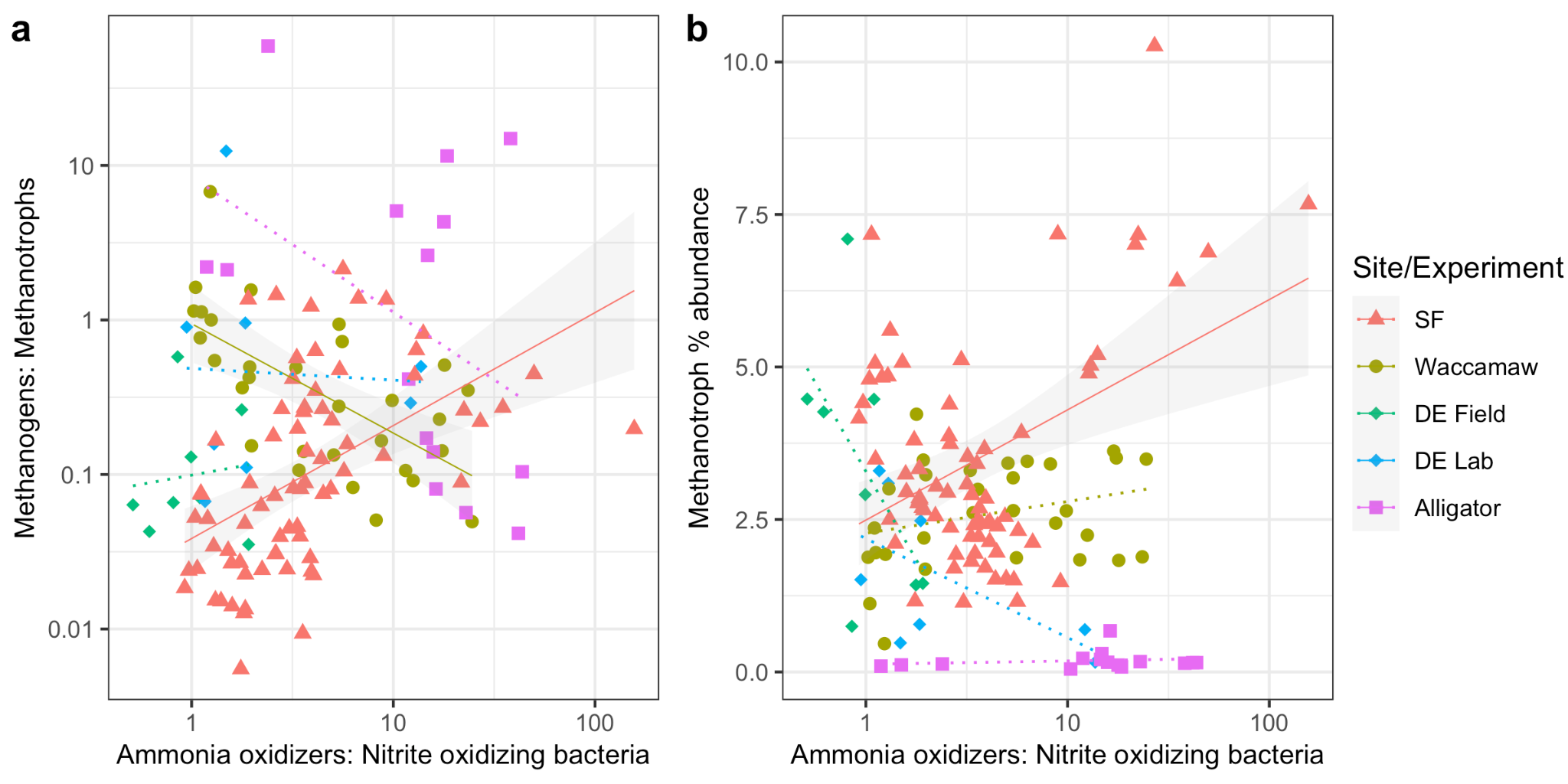

Figure S5. Relationships between ammonia oxidizers (AOA + AOB): nitrite oxidizing bacteria ratios and (a) methanogens: methanotrophs ratio and (b) methanotroph percent relative abundances. Solid lines represent significant linear regression lines while dotted lines represent non-significant linear regression lines.

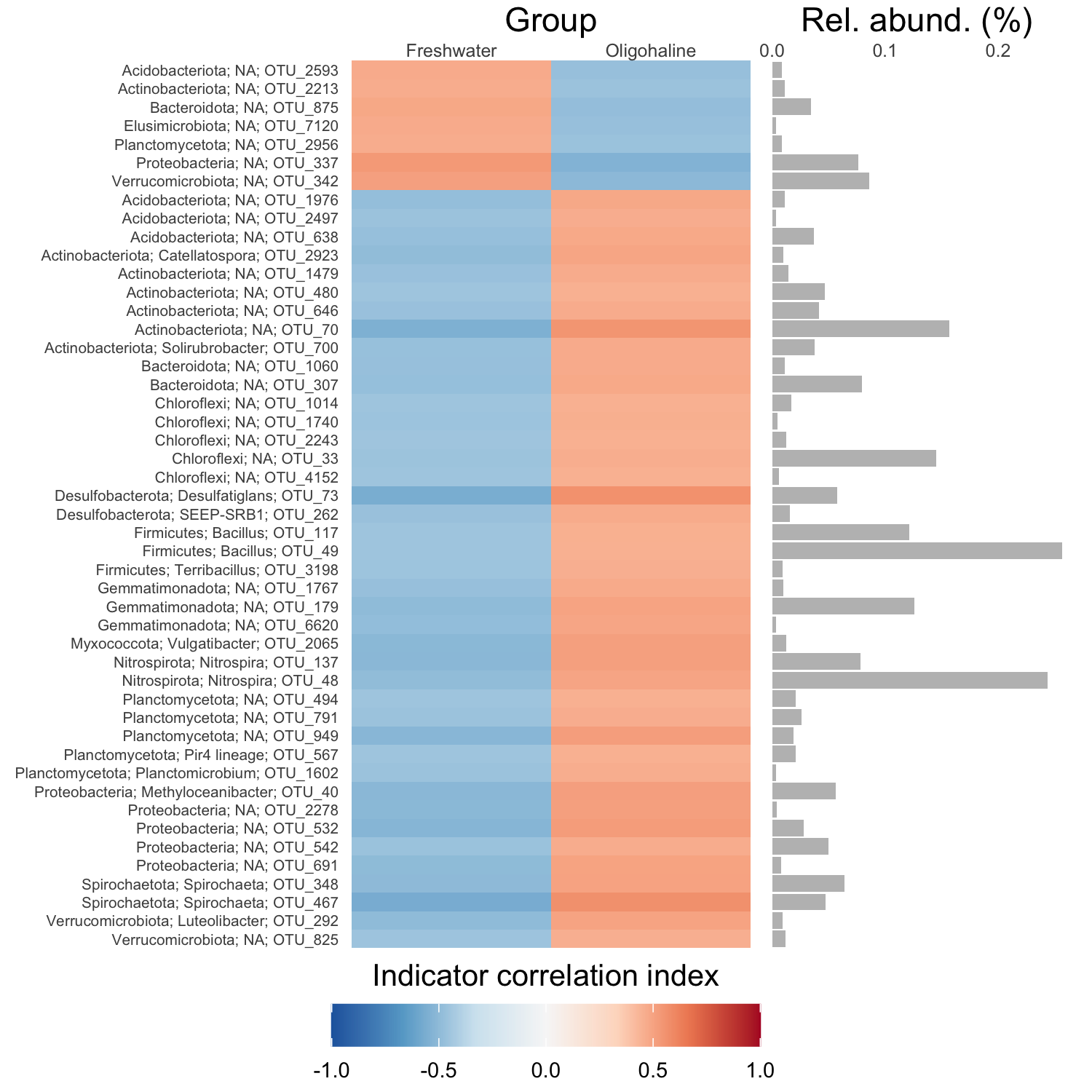

Figure S6. Indicator taxa of samples with freshwater and oligohaline salinities, and their relative abundances. The analysis was conducted with the whole dataset. Only taxa with r > 0.5 are shown here.

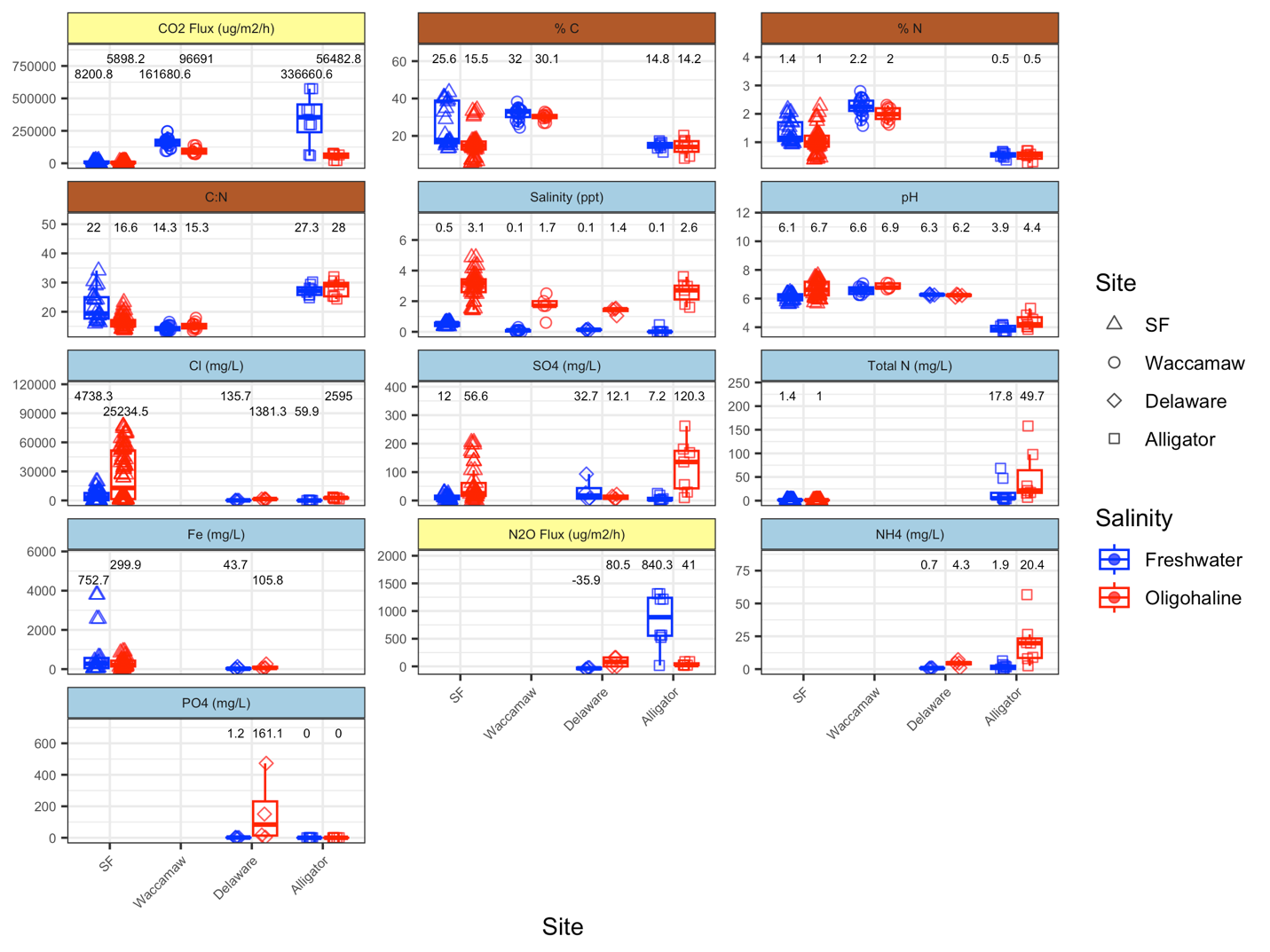

Figure S7. Biogeochemical variables measured in at least two sites. Delaware here refers to just the Delaware field transplant experiment, except for pH, which was measured only in the Delaware lab experiment. Strip colors match those in Figure 5 and Figure S2, with yellow indicating flux data, brown indicating soil data, and blue indicating porewater data. Y-axis units vary among panels and are indicated in the panel titles. Text annotations represent mean values rounded to the nearest tenth; text was repelled vertically for legibility and text location on the y-axis does not necessarily correspond to relative differences in values.

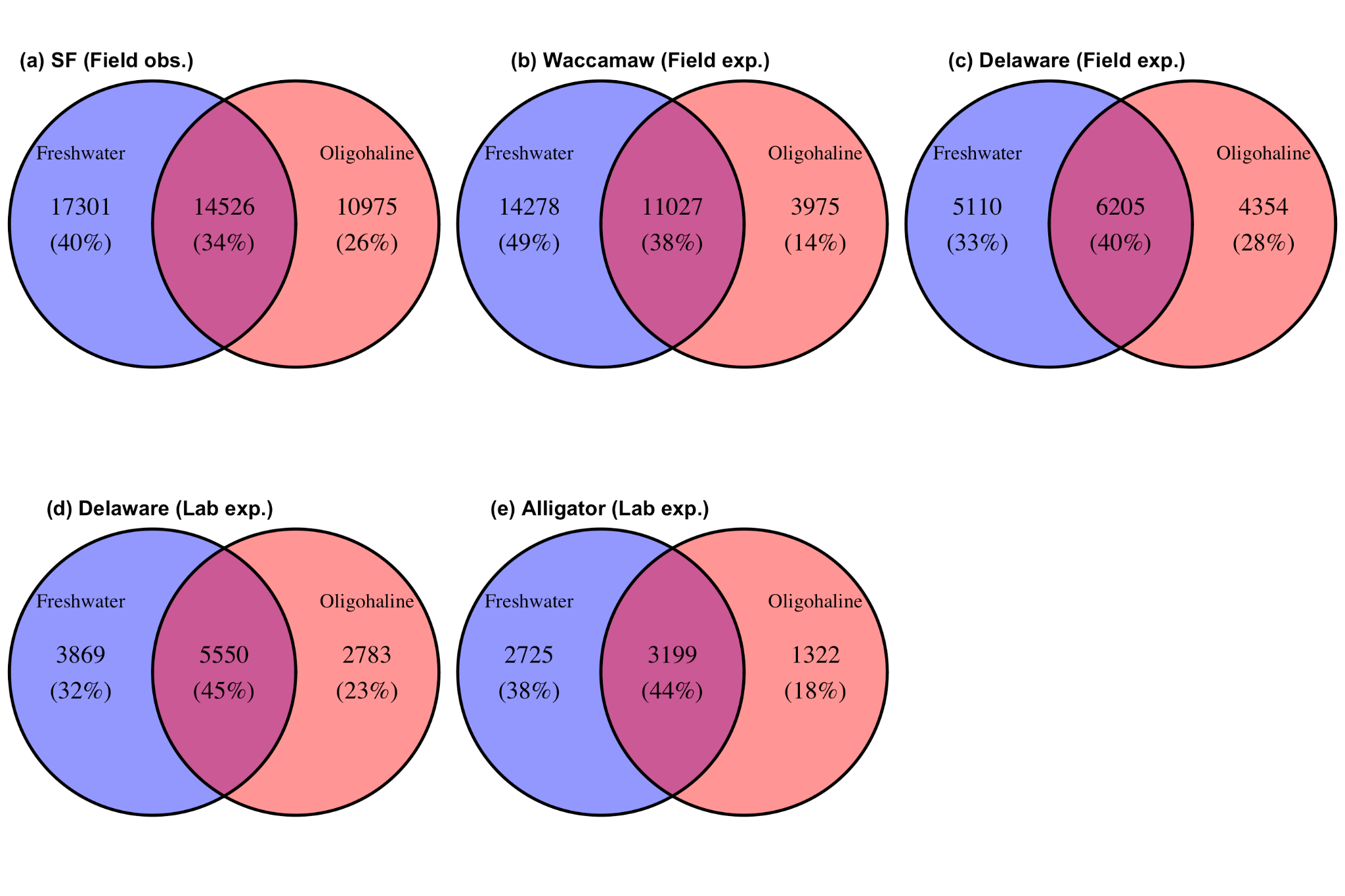

Figure S8. Venn diagrams showing the number of OTUs unique to freshwater samples, unique to oligohaline samples, or shared among the two salinity classes for each site and experiment.

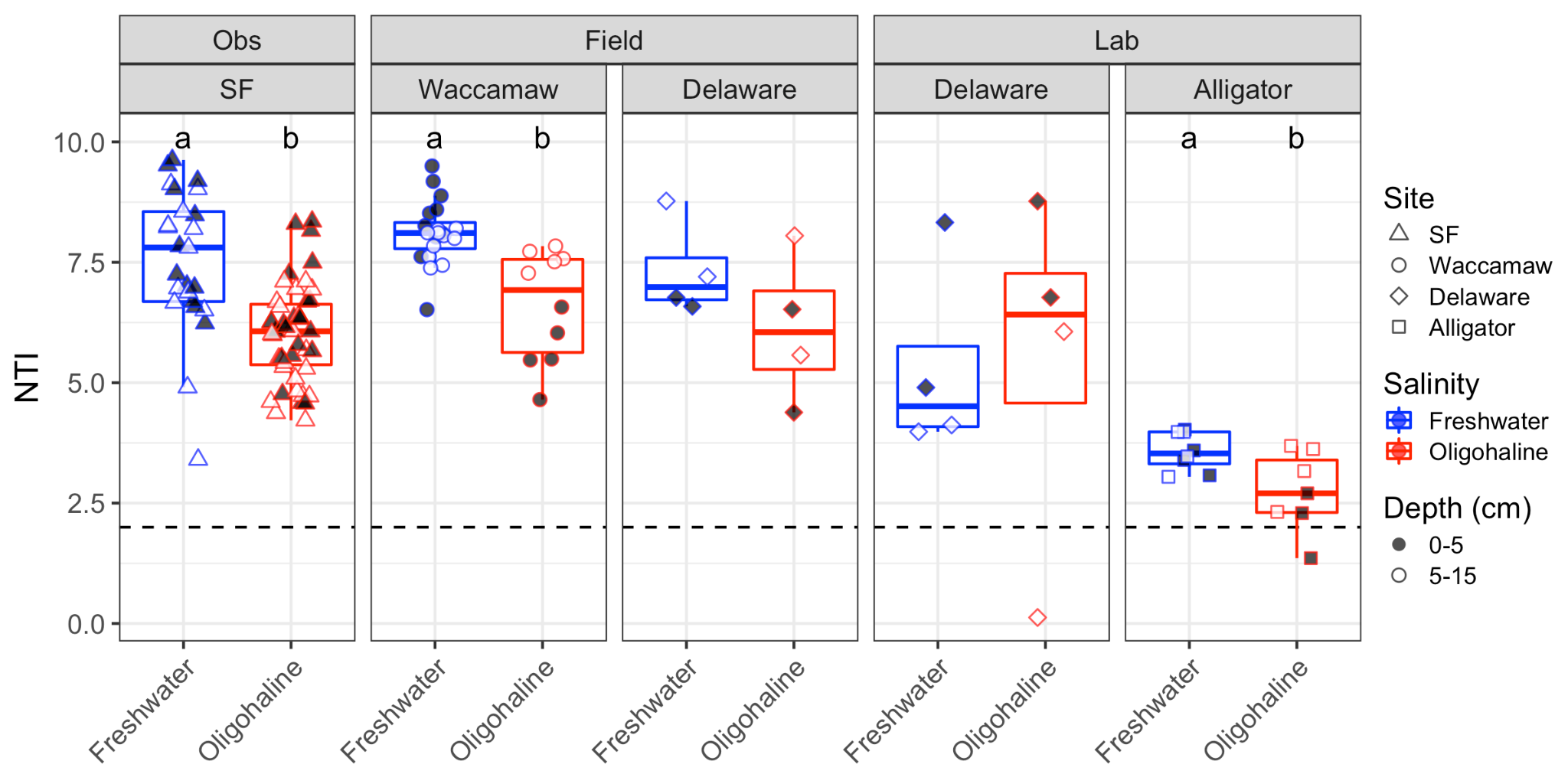

Figure S9. Nearest taxon index (NTI) for each sample, grouped by salinity class and site/experiment and shaded by depth. A dashed line at 2 delineates the boundary between dominance of deterministic processes (> 2) and stochastic processes (< 2 and > -2). Values > 2 indicate phylogenetic clustering.

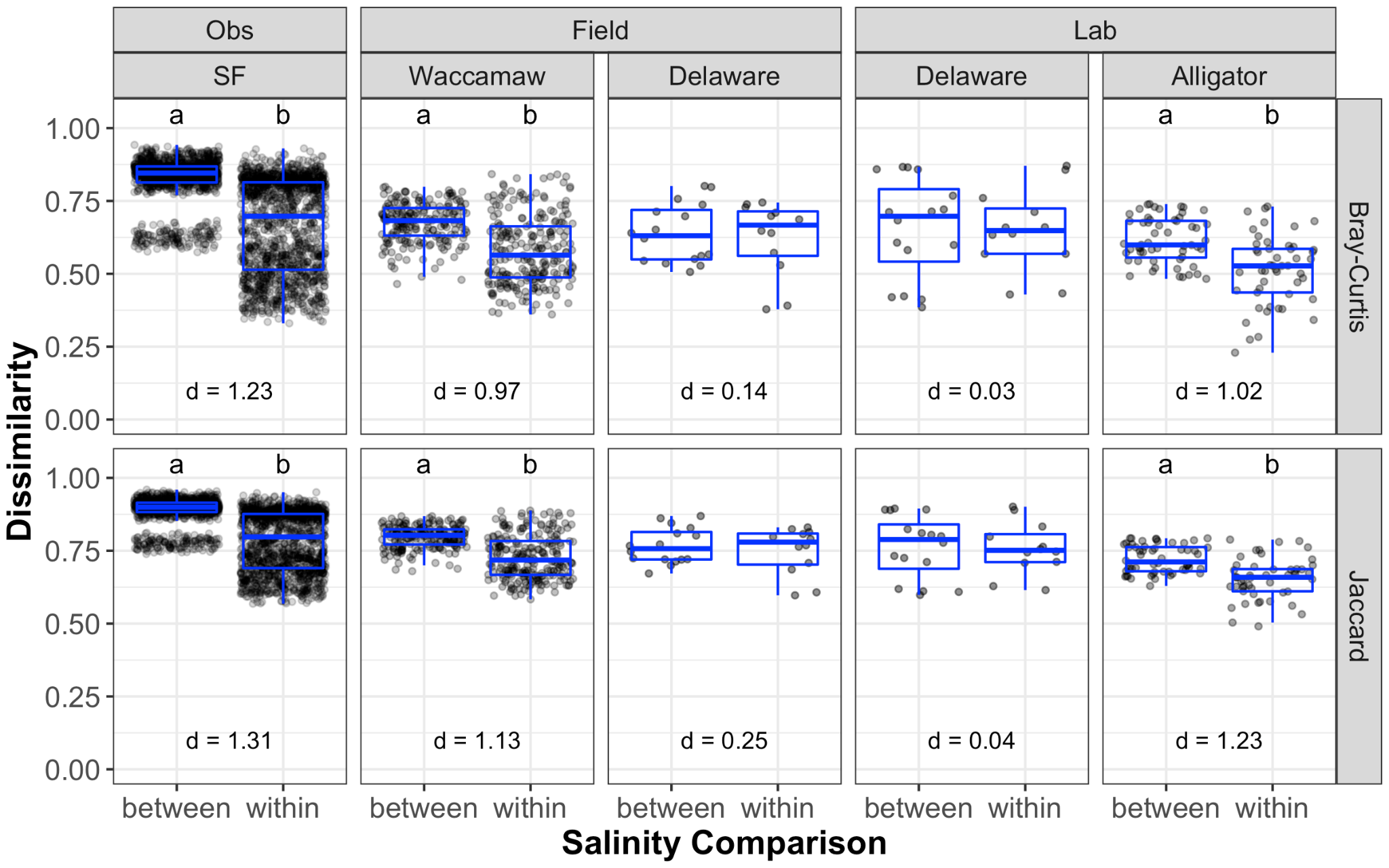

Figure S10. Bray-Curtis dissimilarity and Jaccard dissimilarity for each site/experiment for comparisons between salinity classes (freshwater-oligohaline) or within salinity classes (both freshwater-freshwater and oligohaline-oligohaline comparisons). Different letters represent significant differences. Also shown are Cohen’s *d* effect size values, with larger values indicating a greater difference in dissimilarity between the “between” and “within” comparisons.

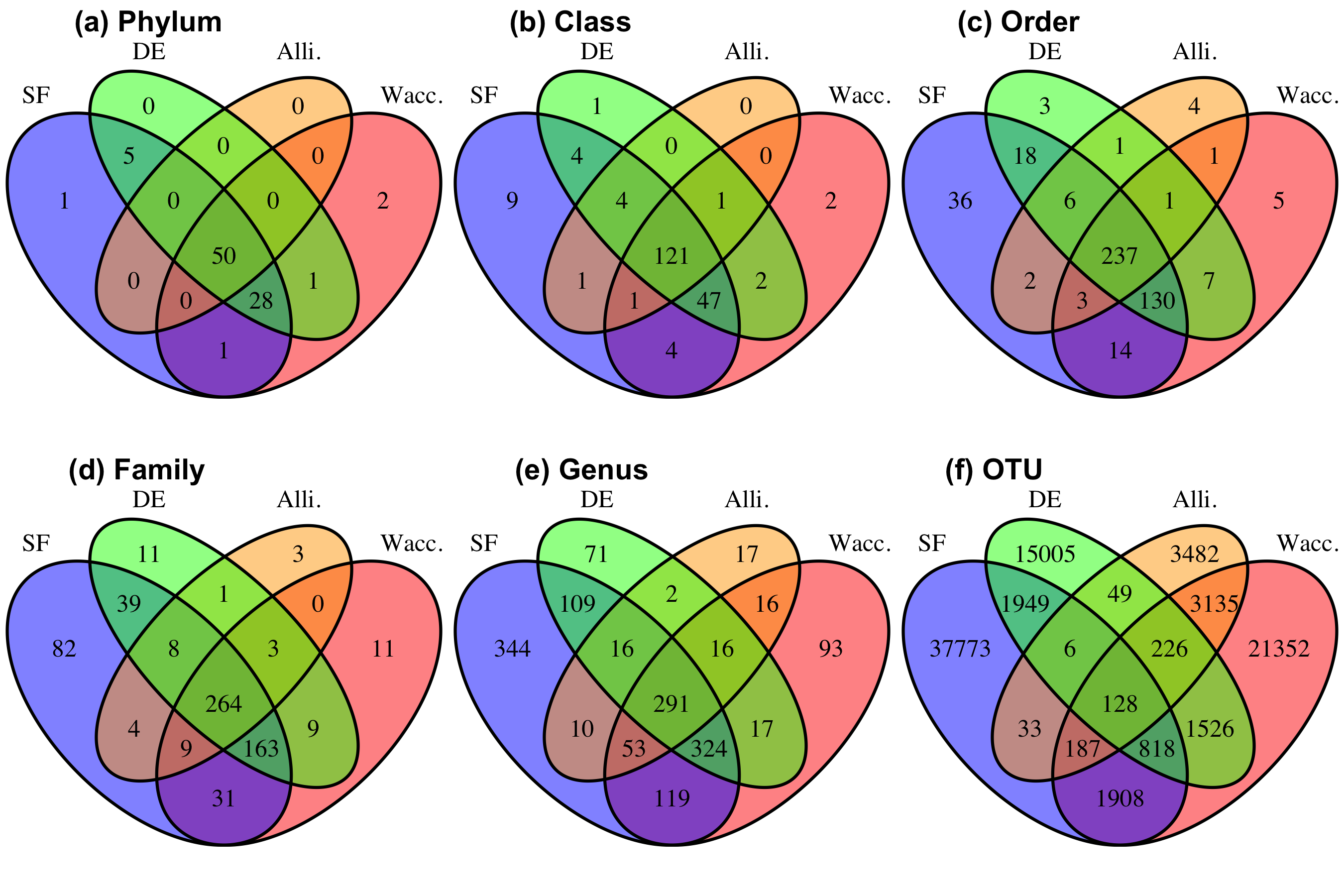

Figure S11. Venn diagrams showing the number of different taxa unique to sites or shared among different combinations of sites. SF = San Francisco Bay and Delta, California, DE = Delaware River, New Jersey, Alli. = Alligator River, North Carolina, Wacc. = Waccamaw River, South Carolina. In this figure DE is the combined laboratory and field experiments from DE.
